# Enhanced Protein-Protein Interaction Discovery via AlphaFold-Multimer

**DOI:** 10.1101/2024.02.19.580970

**Authors:** Ah-Ram Kim, Yanhui Hu, Aram Comjean, Jonathan Rodiger, Stephanie E. Mohr, Norbert Perrimon

**Affiliations:** Department of Genetics, Blavatnik Institute, Harvard Medical School, Boston, Massachusetts, USA; Howard Hughes Medical Institute, Boston, Massachusetts, USA

## Abstract

Accurately mapping protein-protein interactions (PPIs) is critical for elucidating cellular functions and has significant implications for health and disease. Conventional experimental approaches, while foundational, often fall short in capturing direct, dynamic interactions, especially those with transient or small interfaces. Our study leverages AlphaFold-Multimer (AFM) to re-evaluate high-confidence PPI datasets from *Drosophila* and human. Our analysis uncovers a significant limitation of the AFM-derived interface pTM (ipTM) metric, which, while reflective of structural integrity, can miss physiologically relevant interactions at small interfaces or within flexible regions. To bridge this gap, we introduce the Local Interaction Score (LIS), derived from AFM’s Predicted Aligned Error (PAE), focusing on areas with low PAE values, indicative of the high confidence in interaction predictions. The LIS method demonstrates enhanced sensitivity in detecting PPIs, particularly among those that involve flexible and small interfaces. By applying LIS to large-scale *Drosophila* datasets, we enhance the detection of direct interactions. Moreover, we present FlyPredictome, an online platform that integrates our AFM-based predictions with additional information such as gene expression correlations and subcellular localization predictions. This study not only improves upon AFM’s utility in PPI prediction but also highlights the potential of computational methods to complement and enhance experimental approaches in the identification of PPI networks.

## Introduction

Protein-protein interactions (PPIs) play pivotal roles in nearly all biological functions, regulating a wide range of cellular processes, from controlling intracellular enzymatic activities to sensing extracellular signals. Alterations or disruptions in PPIs can lead to a variety of diseases, including cancers and metabolic syndromes. Moreover, the importance of PPIs extends far beyond academic interest; identifying PPIs is crucial for advancing diagnostics and therapeutic interventions [1, 2]. As a result, high-confidence identification of PPIs has long been a priority for the biological and biomedical communities.

Over the decades, the identification of PPIs at proteome scale has primarily relied on experimental approaches such as immunoprecipitation-mass spectrometry (IP-MS) and yeast two-hybrid screening (Y2H) [3–11]. Despite the contributions made by application of these experimental approaches to identifying PPIs, they come with inherent challenges. For example, false negative discovery in Y2H necessitates the application of multiple experimental assay versions [12], and the interactions identified using IP-MS are not necessarily binary in nature. Consequently, following a large-scale study, follow-up assays are often needed to address false discovery issues and identify the subset of PPIs that represent true direct interactors. These supplementary approaches also present challenges and are impractical to apply at large scale.

Recent innovations in protein structure prediction methods such as development of AlphaFold (AF) [13, 14] and RoseTTAFold [15] herald a transformative phase in the field of protein structure biology, as these methods can be used to predict protein structures with remarkable accuracy [16]. The more recently developed AlphaFold-Multimer (AFM) approach has further expanded these capabilities by predicting interactions between pairs of proteins [17]. AFM has catalyzed a significant shift in the landscape of PPI predictions [18], leading to breakthroughs across various organisms and protein types [19–27]. AFM not only offers an innovative and scalable alternative to experimental methods such as IP-MS and Y2H but also can be used to help address their inherent limitations.

Since AFM was launched, several groups have reported methods designed to evaluate [28–30], extend [31–33], and improve [34–39] the performance of AFM in general or for specific applications. Metrics reflecting the predicted structural accuracy have often been used to distinguish between positive and negative PPIs [19, 24, 28, 31, 40, 41]. However, inaccuracies of computational approaches used to predict PPIs might result from difficulties in the underlying protein structure predictions. Furthermore, many physiologically relevant interactions depend on highly localized, small interfaces, including intrinsically disordered regions (IDRs) [42–46].

Flexible regions like IDRs often decrease overall protein structure accuracy. Therefore, structural accuracy-based metrics might not be sufficient, especially when predicting interactions involving proteins with flexible conformations. This highlights the need for the development of new measures capable of detecting direct interactions, particularly in cases involving localized or variable protein domains, to better encompass the various types of PPIs that we know occur *in vivo*.

In this work, we applied and further adapted AFM to reanalyze high-throughput experimental datasets. Through an analysis of datasets of high-confidence PPIs and matched control sets for *Drosophila* and human, we identified limitations associated with using the interface pTM (ipTM) metric derived from AFM, which is a measure of the structural accuracy of a predicted protein complex, as the basis for PPI identification. To overcome these limitations, we refined the AFM approach by using the Predicted Aligned Error (PAE) data provided in AFM outputs to develop a Local Interaction Score (LIS) that focuses on areas with low PAE. Using our PAE-based LIS approach, we were able to predict direct PPIs within an IP-MS dataset and identify high-confidence PPIs within Y2H datasets with higher sensitivity than what can be achieved with AFM alone.

As a specific test case, we applied the LIS approach to the *Drosophila* m^6^A methyltransferase complex (MTC) IP-MS dataset [47]. Our analysis re-constructed known components of the MTC, and identified both literature-supported and potential new direct interactions. We also extended application of the LIS approach to large-scale *Drosophila* datasets generated by IP-MS [7] or Y2H [11], and lists curated from the literature, leading to the prediction of over 5,000 direct PPIs after the evaluation of more than 30,000 potential interactions. In addition to the evaluation of existing datasets, we also applied the LIS approach for predicting potential new interactions that are related to major signaling pathways, metabolic pathways as well as kinase-substrate relationships. All together, we evaluated more than 80,000 fly protein-pairs with 8643 genes involved and the number will continue to increase with more predictions being processed. Finally, we developed the FlyPredictome, an online resource that makes it possible for researchers to explore AFM-predicted results alongside protein-centric information from other resources such as expression patterns and subcellular localization predictions.

## RESULTS

### Limitations of assessing PPIs using AFM-generated ipTM scores

AFM has significantly advanced structure-based prediction of PPIs. Specifically, the AFM-derived ipTM score has emerged as a standard metric for assessing the structural integrity of predicted complexes. High ipTM scores are often associated with stable, enduring PPIs, as corroborated by experimental structure determinations [17]. Nonetheless, a focus on structural accuracy might not provide a complete representation of the various types of PPIs that take place within physiological environments, where interactions are often local and transient. We reasoned that there is additional information embedded in AFM outputs that could be used to generate high-confidence predictions of PPIs even when the overall ipTM scores are not suggestive of an interaction, thereby improving the sensitivity of AFM-based PPI predictions. Specifically, we predicted that the color-coded plots of PAE that are generated as part of AFM outputs are a potential source of data useful for PPI prediction at improved sensitivity.

To test the potential use of information from PAE plots to predict PPIs, we first identified previously assembled sets of well-supported PPIs, as well as control sets in the form of random or artificially generated lists of proteins pairs. Specifically, we made use of ‘positive reference sets’ (PRSs) and corresponding ‘random reference sets’ (RRSs) previously established for *Drosophila* [11] and human proteins [6]. Using the *Drosophila* and human PRSs, which are based on experimentally observed PPIs, helps avoid a bias in our testing towards very stable interactions and thus, allows us to more accurately evaluate the relevance of ipTM scores for PPIs that exhibit temporal dynamics in living organisms. Moreover, in addition to using the established RRSs as controls, we generated supplemental control sets comprised of most proteins in the *Drosophila* or human PRS paired with either GFP or *Drosophila* Wingless (Wg) (hereafter, the fly or human ‘PRS-GFP’ and ‘PRS-Wg’ sets). GFP and Wg were selected for their low likelihood of engaging in PPIs with PRS proteins. GFP is widely used as a reporter due to its non-disruptive nature in various species and Wg is a secreted protein expected to have a very specific and limited number of interactions. Thus, the PRS-GFP and PRS-Wg sets serve as additional negative controls.

We first evaluated the PRSs and control sets using the standard AFM method. For each interaction, we generated five separate models, each subjected to five cycles of refinement, to enhance the accuracy of our predictions. The average scores derived from these five models are referred to as the ‘average ipTM,’ and the score from the single best-performing model determined by AFM is designated as the ‘best ipTM.’ As anticipated, both the fly and human PRSs exhibited higher average and best ipTM scores compared to the corresponding fly and human RRS, PRS-GFP, and PRS-Wg pairs (Figures 1B-E, Supplementary Figure 1). Furthermore, when averaged, ipTM scores for PRS pairs were significantly different from control ipTM scores (p-values: 1.38e-175 for average, 8.60e-122 for best) (Figure 1F-G). These indicate that ipTM-based differentiation between the PRSs and control sets is indeed feasible.

**Figure 1.**
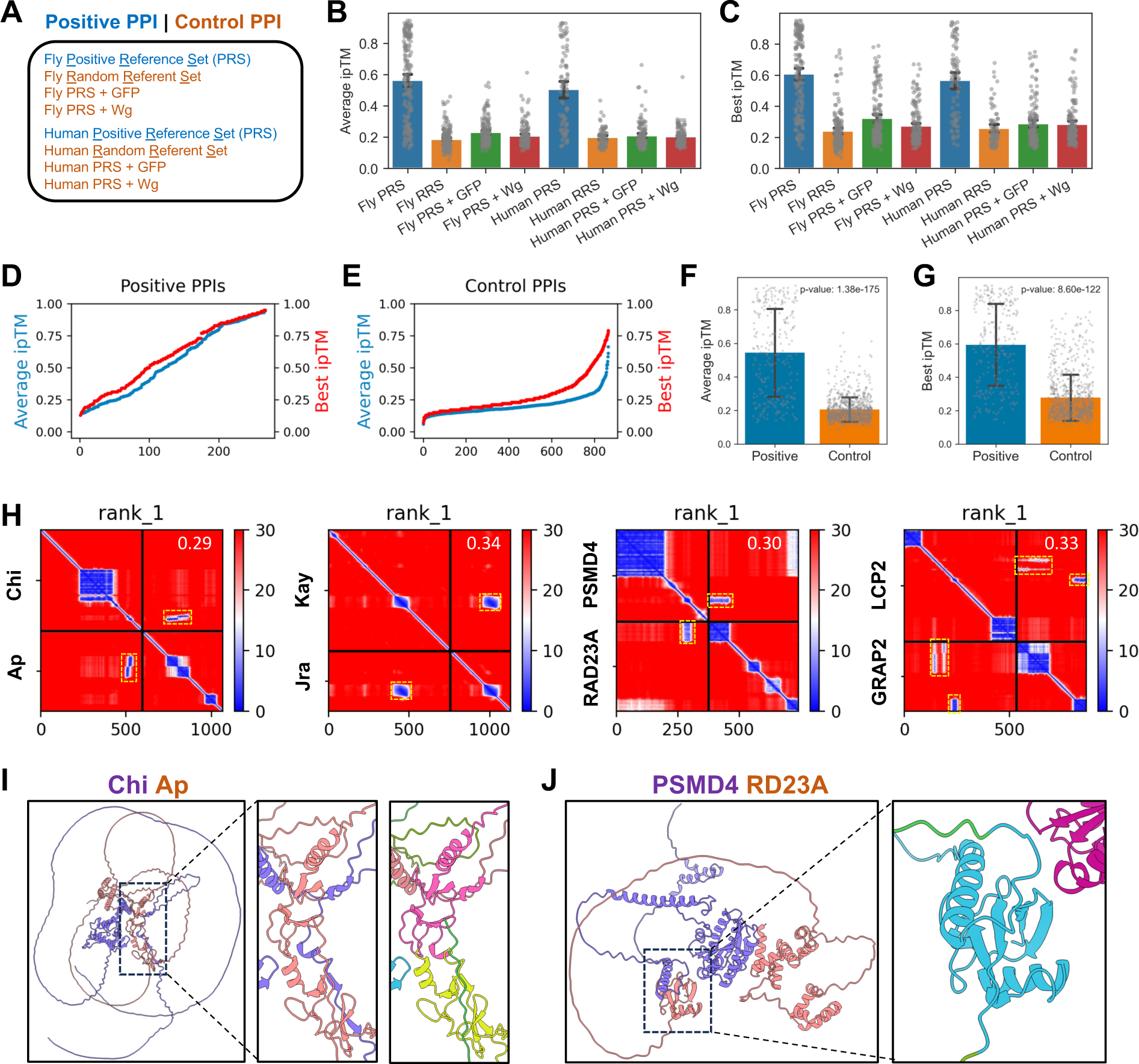
Evaluation of ipTM scores across *Drosophila* and human protein-protein interaction sets. A. Overview of protein-protein interaction (PPI) test sets. Positive PPIs comprise *Drosophila* and human Positive Reference Sets (PRS), while negative PPIs include published *Drosophila* and human Random Reference Sets (RRS), as well as the combinations of human and fly PRS with GFP (PRS + GFP) or *Drosophila* Wingless (Wg) protein (PRS + Wg). B. Mean average of the ipTM scores across test sets, showing positive PPI sets (fly and human PRS) have higher mean average ipTM than individual control PPI sets. C. Mean best ipTM scores across test sets, similarly indicating higher values for positive PPI sets compared to individual control sets. D. Cumulative distribution of the average ipTM scores and the best ipTM scores for positive PPI sets, with the blue color representing the average ipTM scores while the red color representing the best ipTM scores, respectively. The x-axis indicates the number of PPIs. E. Cumulative distribution of the average ipTM scores and the best ipTM scores for control PPI sets, with the blue color representing the average ipTM scores while the red color representing the best ipTM scores, respectively. The x-axis indicates the number of PPIs. F. The comparison of the mean average ipTM scores for positive PPI sets versus control PPI sets, with statistical significance indicated by a p-value from t-test. G. The comparison of the mean best ipTM scores for positive PPI sets versus control PPI sets, with statistical significance indicated by a p-value from t-test. H. Examples of the low ipTM-score predictions with discernible local interactions in PAE maps, outlined by yellow dotted boxes. The four pairs for the interactions involved are *Drosophila* Chi vs Ap, *Drosophila* Kay vs Jra, human PSMD4 vs RAD23A, and human LCP2 vs GRAP2 with the ipTM scores shown in the top right corner. I. 3D view of the predicted complex structure of *Drosophila* Chi and Ap proteins. Interacting domains are highlighted in pink and yellow-green, as determined using ChimeraX domain analysis. J. 3D view of the predicted complex structure of human PSMD4 and RD23A proteins. The interacting domain is highlighted in cyan, showcasing the domain despite the low structural accuracy.

However, we noticed that a few PPIs with low ipTM scores nevertheless had significant areas of blue color on PAE maps, indicative of regions with lower alignment error that we interpret to be probable sites of interaction between the two proteins (Figure 1H, Supplementary Figure 2). The predicted structures of *Drosophila* Chi-Ap [48, 49] and human PSMD4-RD23A [50–52] exemplify this observation. For these pairs, the local interaction domains have low PAE values despite the overall low ipTM scores assigned by AFM to these protein pairs (Figure 1I-J). These cases typically involve proteins with predicted flexible regions and for which interactions were localized to specific sub-domains of one or both proteins. Based on this, we predicted that information in the PAE maps could be used as a supplement or alternative to ipTM to improve the sensitivity of PPI predictions.

### Enhancing PPI detection using Local Interaction Score (LIS)

AFM analyses provide PAE maps that align the interaction sites of each protein pair at the level of individual amino acids, with PAE values color-coded in a gradient from red to blue. The darkest blue color indicates the lowest PAE values, hence indicating the highest confidence in an interaction prediction. To exploit this feature, we first selected the areas of blue color, which we refer to as Local Interaction Areas (LIAs), while ignoring areas with higher PAE values (in red), which are unlikely to correspond to interaction zones (Figure 2A). By inversely mapping PAE values within the LIA, we derived a value, ranging from 0 to 1, for which a higher number denotes a stronger inferred interaction. We then calculate a Local Interaction Score (LIS) by averaging the inverted PAE values across the interfaces of the interacting protein pair. We tested the sensitivity and specificity of this approach using the PRSs and control sets, comparing results obtained using PAE values from 1 to 30 as the cutoff. From this analysis, we found that a PAE cutoff of 12 maximizes the area under the curve (AUC) in a receiver operating characteristic (ROC) analysis, yielding an average LIS of 0.911 and a best LIS value of 0.891 (Supplementary Figure 3). These values indicate the highest level of discriminatory power when differentiating between positive and negative interactions in our testing sets. A PAE cutoff of 12 was therefore established as our standard for further analysis.

**Figure 2.**
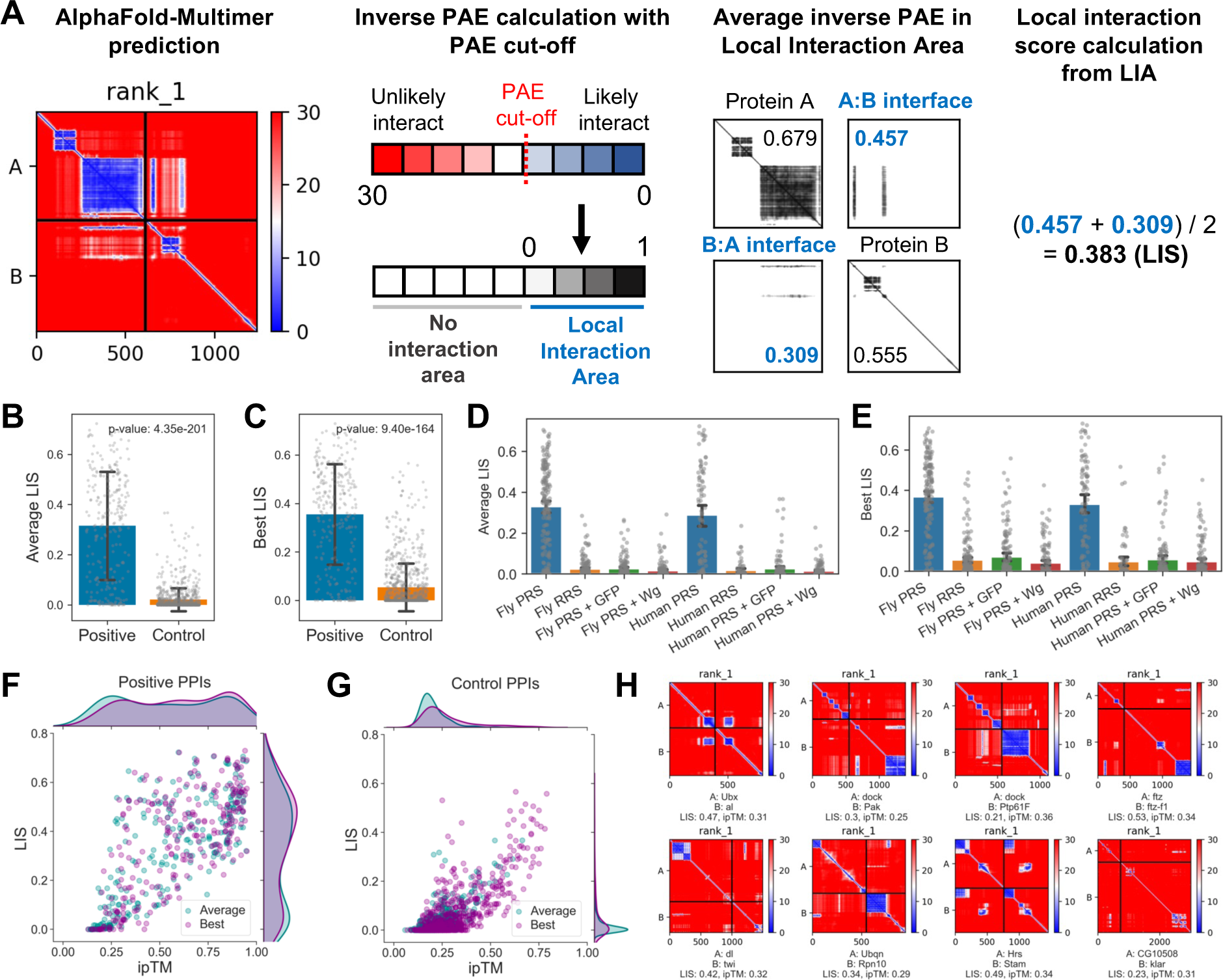
Development and validation of the Local Interaction Score for identifying PPIs. A. Illustration of the algorithm for identifying the Local Interaction Area (LIA) and calculating the Local Interaction Score (LIS). Following AlphaFold-Multimer predictions, Predicted Aligned Error (PAE) values, indicative of individual amino acid interaction likelihood, are extracted from ColabFold outputs. A PAE cutoff is applied, ignoring amino acid contacts with higher PAE values while retaining those below the threshold as part of the LIA. PAE values within the LIA are inverted to a scale of 0 to 1, where higher values denote stronger predicted interactions. The average of these inverted PAE values at the interaction interfaces yields the LIS. B. The comparison of the mean average LIS for the positive PPI set versus control PPI set, with the p-value from the t-test. C. The comparison of the mean best LIS for the positive PPI set versus control PPI set, with the p-value from the t-test. D. Bar graph depicting the mean average LIS across individual PPI sets. E. Bar graph depicting the mean best LIS across individual PPI sets. F. Dot plot to show the relationship between the ipTM and the LIS for positive PPI sets, where the average ipTM/LIS are represented as blue dots and the best ipTM/LIS as purple dots. G. Dot plot to show the relationship between the ipTM and the LIS for control PPI sets, where the average ipTM/LIS are represented as blue dots and the best ipTM/LIS as purple dots. H. Example PAE maps from direct *Drosophila* PPI predictions showing areas of interaction (blue) despite low overall ipTM scores, with the specific LIS and ipTM values annotated for each case.

In our analysis, we found that the PRSs had significantly higher average LIS values (p-value: 4.35e-201) and best LIS values (p-value: 9.40e-164) as compared to the control sets (Figure 2B-C), indicating a robust separation. This separation was greater than that observed with ipTM scores (p-values: 1.38e-175 for average ipTM and 8.60e-122 for best ipTM) (Figure 1B-C). Detailed individual test set results using average and best LISs, as depicted in Figure 2D-E and Supplementary Figure 4, show a clear distinction between PRSs and control sets.

Visualization of LISs and ipTM scores for the PRSs and control sets reveals that LISs and ipTM scores correlate well in general while identifying many PPIs from PRS with low ipTM scores but relatively high LIS values (Figure 2F-G). This indicates that there are interactions that ipTM-based evaluations could miss, as exemplified in Figure 2H. Thus, our results suggest that physiologically relevant PPIs, especially those characterized by localized and flexible interactions that can be missed using ipTM-based evaluation, can be identified and assessed using a PAE-based metric such as the LIS.

### Comparison of the LIS with existing AFM-derived metrics

We next compared results obtained with the LIS approach to results obtained using other metrics, such as Model Confidence (0.2*ipTM + 0.8*pTM), which is embedded in AFM [17]; pDockQ [24, 53] and pDockQ2 [29]; and the LIA values we calculate prior to LIS derivation. ROC analyses of the same PPI datasets revealed that the average LIS scored the highest AUC, followed by average ipTM, and then average LIA (Figure 3A). Among ‘best’ metrics, best LIS was the leading metric in terms of AUC, followed by best LIA (Figure 3B). This indicates that both the average and best LIS are effective in discerning positive PPIs within our test sets. The results of analysis with each of the approaches we compared are shown in Supplementary Figure 5, Supplementary Figure 6, and Supplementary Figure 7.

**Figure 3.**
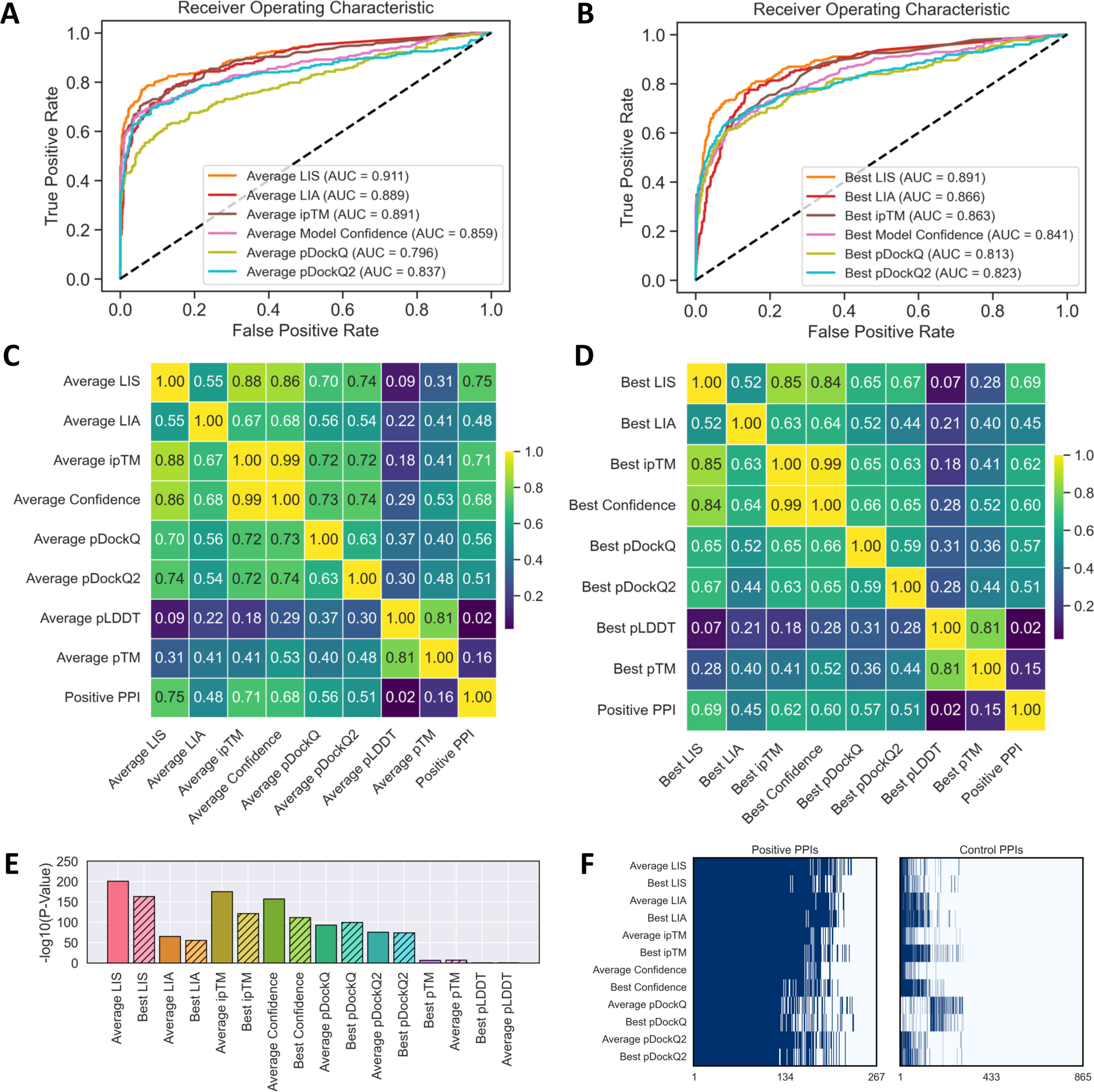
Comparative analysis of LIS with established AlphaFold-Multimer metrics. A. Receiver operating characteristic (ROC) analysis for average score metrics comparing LIS and other AFM metrics. True positives are based on fly and human PRSs, and false positives are based on control PPI sets. The AUC for each metric is provided, illustrating that LIS demonstrates best performance. B. ROC analysis for the best score metrics. AUC values indicate the best performance of LIS in identifying direct interactions. C. Correlation heatmap among the average score metrics. The heatmap illustrates the correlation of each metric with positive PPIs and with structural accuracy indicators pLDDT and pTM. LIS shows the highest correlation with positive PPIs and the lowest with pLDDT and pTM, suggesting its higher sensitivity and independence from structural accuracy. D. Correlation heatmap among the best score metrics. This heatmap also demonstrates the correlation of each metric with positive PPIs and accuracy indicators. LIS maintains the highest correlation with positive PPIs and the lowest with structural accuracy, confirming its utility. E. P-value comparison using various metrics between positive and control PPI sets. Both average and best LIS metrics show the most statistically significant differences, with the lowest p-values indicating robust discrimination. F. Heatmap of PPI predictions using optimal thresholds from ROC analysis. PPIs exceeding the optimal threshold for each metric are identified as direct interactions and highlighted in blue.

The optimal thresholds derived from ROC analysis for these metrics, as well as their corresponding sensitivity, specificity, and Youden’s Index, are presented in the Supplementary Figure 8. LIS consistently showed strong performance in nearly all comparisons. Correlation heatmaps for both average metrics (Figure 3C) and best metrics (Figure 3D) aligned well with positive PPIs, particularly for average and best LIS. To assess whether each metric is influenced by prediction accuracy, we examined their correlations with pLDDT, an indicator of individual amino acid accuracy, and pTM, which represents predicted monomer structure accuracy (Figure 3C-D, Supplementary Figure 9). Among all the metrics, LIS exhibited the lowest correlation with pLDDT, which indicates individual amino acid accuracy, and pTM, representing predicted monomer structure accuracy. This suggests that LIS has the capability to identify potential PPIs regardless of structural accuracy.

When consolidating the metrics across PRSs and control sets, average LIS showed the most significant distinction, and best LIS similarly excelled among the best metrics (Figure 3E). This highlights the potential of LIS to effectively discriminate positive PPIs. Utilizing the optimal thresholds determined by ROC analysis (Figure 3A-B, Supplementary Figure 8), we categorized interactions as ‘supported’ (positive) or ‘unsupported’ (negative) PPIs. Figure 3F and Supplementary Figure 10 illustrate that curated positive PPI sets (i.e., the fly PRS and human PRS) consistently showed a greater prevalence of positive PPIs in comparison to control sets in all tested metrics.

### Predicting direct interactions among yeast PPIs and Eukaryotic Linear Motif resource

Having established that LIS provides a new and valid way to predict PPIs with small interaction interfaces, we next expanded our analysis to a *Saccharomyces cerevisiae* PRS and RRS [4], and to pairs in the Eukaryotic Linear Motif (ELM) resource [46, 54], which contains experimentally determined protein complex structures involving short linear motifs. We hypothesized that these additional datasets would likely include PPIs that rely on small interfaces. Using a similar approach as our previous simulations, we generated yeast PRS-GFP and yeast PRS-Wg sets as additional negative control sets. As shown in Figure 4A-D, yeast PRS pairs and ELM-annotated PPIs resulted in high ‘average’ and ‘best’ scores in both LIS and ipTM analyses as compared with the yeast RRS, yeast PRS-GFP, and PRS-Wg.

**Figure 4.**
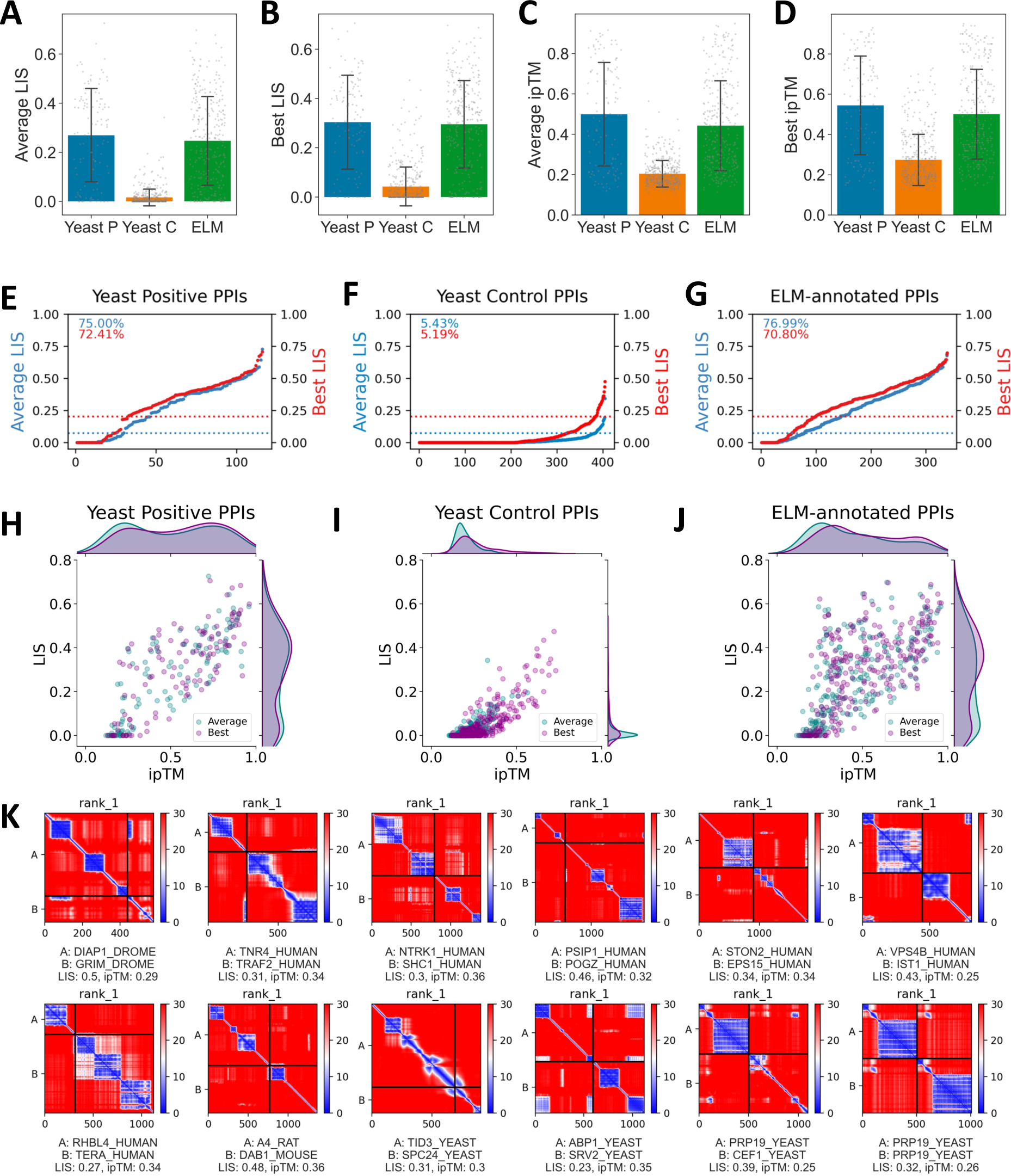
Assessment of LIS and ipTM in yeast PPI sets and ELM-annotated PPIs. A. Comparison of the mean average LIS across three groups: yeast positive PPIs (Yeast P), yeast control PPI sets (Yeast C), and ELM-annotated PPIs (ELM). B. Comparison of the mean best LIS across the same three groups as in (A). C. Comparison of the mean average ipTM scores across the same three groups as in (A), highlighting differences in structural accuracy predictions. D. Comparison of the mean best ipTM scores for the same three groups as in (A). E. Cumulative distribution of average and best LIS for yeast positive PPIs. Percentages at the top of each graph indicate the proportion of PPIs exceeding optimal LIS thresholds indicated by dotted lines, and the x-axis indicates the number of PPIs (E-G). F. Cumulative distribution of average and best LIS for yeast control PPIs. G. Cumulative distribution of average and best LIS for ELM-annotated PPIs. H. Scatter plot showing ipTM and LIS for yeast positive PPIs. Average metrics are colored by blue, and best metrics are colored by purple. I. Scatter plot correlating ipTM with LIS for yeast control PPIs, where average metrics are denoted in blue and best metrics in purple (I-J). J. Scatter plot correlating ipTM with LIS for ELM-annotated PPIs. K. PAE maps displaying examples of PPIs with small interaction interfaces from yeast positive and ELM-annotated PPIs. The selected PPIs exhibit best LIS values exceeding the optimal threshold (0.21) and best ipTM scores falling below the optimal threshold (0.38). Names of individual proteins, species, LIS, and ipTM scores are specified for each example.

Applying optimal thresholds for average and best LIS to the yeast PRS, yeast negative control sets, and ELM-annotated PPIs (Figure 4E-G), we observed that over 70% of yeast positive PPIs and ELM-annotated PPIs were predicted to be direct interactions. Approximately 5% of yeast negative control sets had values that exceeded these thresholds, a trend also found in negative sets from *Drosophila* and human. The scatter plots for LIS and ipTM for each set are shown in Figure 4H-J. Compared to yeast negative control sets, both yeast PRS and ELM-annotated PPIs generally showed high LIS and high ipTM scores. However, a substantial proportion of PPIs in ELM-annotated PPIs had low ipTM scores but high LISs, consistent with the idea that these PPIs depend on linear motifs for interaction.

We provide examples of PPIs with low structure accuracy but high LISs in Figure 4K. We identified such PPIs within both the yeast PRS and ELM-annotated PPIs. Notably, when full-length sequences were used in the analyses, even experimentally confirmed PPIs that involve protein-linear peptide interactions did not consistently have high ipTM scores. This is again consistent with the idea that ipTM and structural accuracy-based metrics have limitations in distinguishing between direct and indirect interactions when a PPI relies on a local interface. Overall, these results support the use of LIS analysis to identify PPIs that rely on a small interface.

### Expanding the network from an IP-MS dataset via iterative AFM screening

Previously, we characterized the interactome of m^6^A methyltransferase complex (MTC) through IP-MS, revealing the role of the Mechanistic Target of Rapamycin Complex 1 (mTORC1) in regulating autophagy through m^6^A RNA methylation [47]. While we confirmed several interactions within the complex using co-immunoprecipitation and pull-down assays, the full complexity of the interactome’s internal dynamics required further characterization. Thus, leveraging the LIS to discern direct interactions, we applied LIS analysis alongside sequential AFM prediction to refine and expand the MTC interactome (Figure 5A-B). We observed that some PPIs with very high LIS had very low LIA values, which might be indicative of false positive discovery. To mitigate this, we introduced LIA thresholds as additional filters to distinguish positive and negative PPIs. We initiated this multistep analysis by using both LIS and LIA to filter for direct interactions, considering both average and best scores. If either the average or best scores met the criteria, we considered them indicative of a direct interaction.

**Figure 5.**
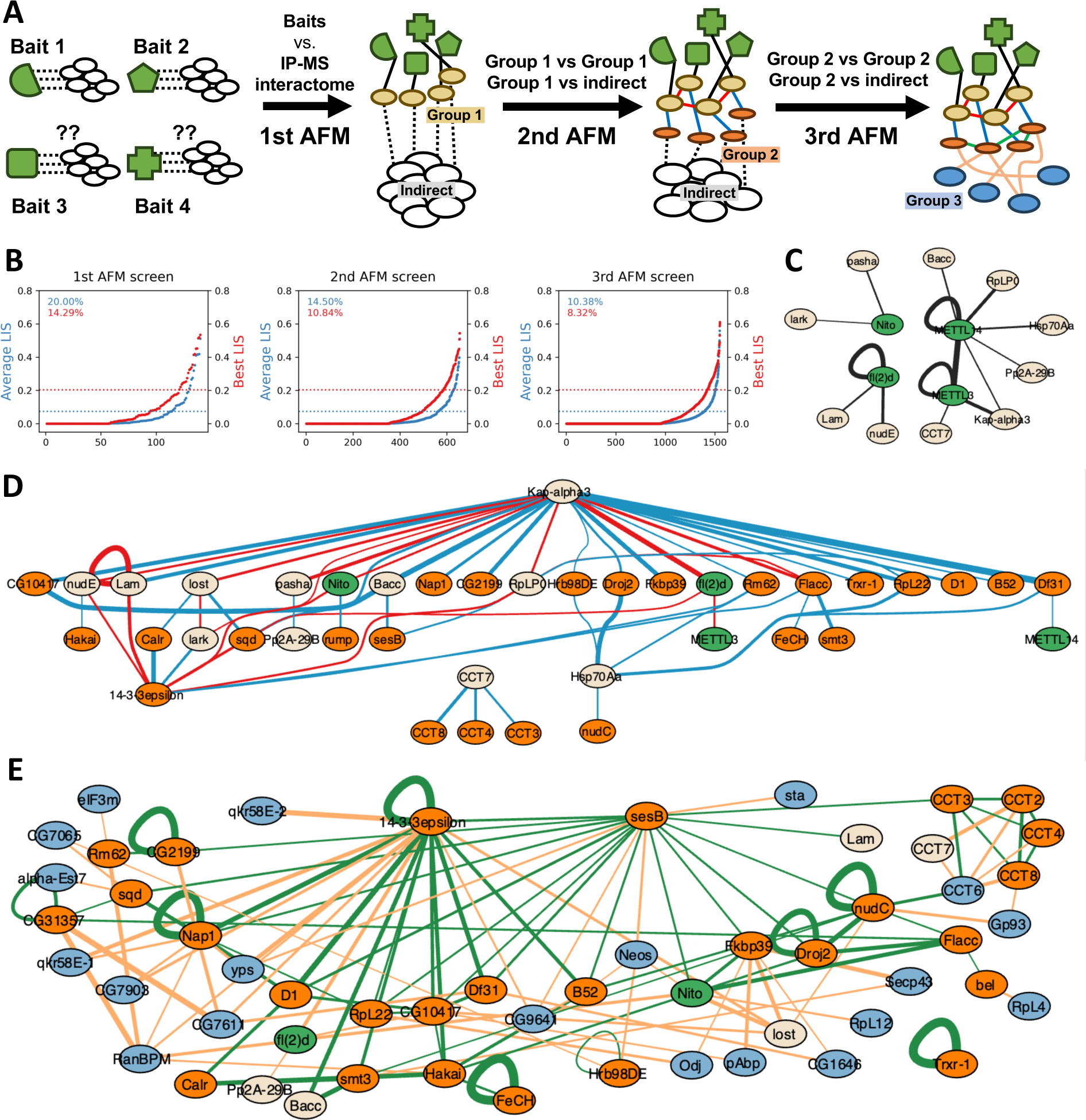
Mapping PPI networks from IP-MS data through iterative AFM screening. A. The scheme shows the steps to build a binary PPI network from an IP-MS dataset using AFM. It starts with the evaluation of all the bait-prey interactions using AFM to find proteins (beige nodes) that directly interact with the bait protein(s) (green nodes). Beige nodes make the Group 1. The second AFM step screens for the connections of each beige node from Group 1 with all the other proteins identified from IP-MS to pinpoint more directly binding proteins which are the orange nodes (Group 2). The third AFM can further expand the map, identifying the binary interactions within Group 2 and beyond, contributing towards a comprehensive binary PPI network. B. Distribution of the average and best LIS for each AFM screening with the MTC IP-MS dataset. Percentages at the top of each graph indicate the proportion of PPIs exceeding optimal LIS thresholds indicated by dotted lines. C. The binary interactome generated from the first round of AFM screening, where the baits are marked in green, and the direct interacting proteins identified from this round of screening are in beige (Group 1). The connections determined at this step are illustrated with black edges. The thickness of these lines corresponds to the average LIS, with thicker lines indicating a higher LIS (C-E). Note that three baits are predicted to form homodimers shown with looped lines connecting a protein to itself. D. The expanded interactome from the second round of AFM screening. The direct interacting proteins only identified from this round make Group 2 and are colored in orange. The binary interactions within Group 1 are indicated by red edges while the interactions identified between Group1 and Group 2 are shown by blue edges. E. The comprehensive interactome resulting from the third round of AFM screening. Group 3 proteins are represented by blue nodes. The interactions within Group 2 proteins are indicated by green edges. The connections between Group 2 and Group 3 proteins are shown by orange edges.

The first AFM prediction between each bait and its associated prey pinpointed direct interactions between core proteins and their neighbors (Figure 5C, Supplementary Figure 11A,B), consistent with the well-established observation of heterodimer formation among some baits and specific direct interactions, e.g., binding of METTL3 to METTL14 [55, 56].

The subsequent second AFM prediction focused on proteins from the first group and their interactions with the remaining proteins. This revealed connections within the first group and between the first and second groups (Figure 5D, Supplementary Figure 11C,D). A key finding was that Kap-alpha3, which is required for nuclear import [57–59], appears as a central hub within the MTC interactome. Kap-alpha3 has predicted associations with several other proteins, including both known and potential new interactions, as detailed in the Supplementary Figure 12.

The third AFM prediction aimed to identify interactions within the second group and their potential connections to remaining proteins (Figure 5E, Supplementary Figure 11E,F). The results of this analysis suggest that 14-3-3 epsilon is a central hub, bridging known and previously unidentified interactors (Supplementary Figure 13). The emergence of Kap-alpha3 and 14-3-3 epsilon as central hubs imply that they play significant roles within the interactome, possibly facilitating nuclear import (Kap-alpha3) and serving as a protein scaffold (14-3-3 epsilon). The full predicted interactome is presented in Supplementary Figure 14.

The iterative screening process incrementally added interactions that correspond to known protein complexes. For example, binary interactions within the CCT complex, which is essential for protein folding and cell/organ growth [60–62], were progressively resolved, starting with prediction of CCT7 as a direct interactor with METTL3. Furthermore, subsequent screens identified additional CCT subunits-CCT3, CCT4, CCT8, and later, CCT2 and CCT6, culminating in binary interaction predictions for six of eight CCT subunits within the MTC IP-MS dataset. These predictions are consistent with the suggested role of the CCT complex in folding the m6A complex [47].

Overall, these results show that sequential application of AFM can predict which proteins have direct interactions with IP baits and expand the PPI network. The results not only reaffirm known protein complexes but also suggest that our approach can be used to predict high-confidence PPI networks useful for development of testable new hypotheses.

### Computational analysis of a large-scale *Drosophila* PPI dataset

*Drosophila* has been instrumental in biomedical science, revealing intricate biological mechanisms conserved among species [63–67]. Large-scale proteomics studies utilizing Y2H and IP-MS assays have identified numerous PPIs in *Drosophila* [3, 7, 11, 68–70]. However, large-scale experimental methods for identifying PPIs face challenges, including high false discovery rates and for IP-MS, difficulty in distinguishing direct versus indirect interactions. We reasoned that application of the LIS/LIA computational approach could help identify high-confidence subsets of large-scale experimental *Drosophila* PPI datasets.

The first analysis we performed focused on our recent large-scale Y2H dataset, known as the FlyBi dataset, generated from multiple all-by-all Y2H screens of 10,000 *Drosophila* proteins [11]. This effort led to identification of 8,723 binary interactions spanning 2,939 proteins that met the standards applied in the FlyBi project for high-confidence Y2H interactions. A subset of these PPIs were experimentally validated using the Mammalian Protein-Protein Interaction Trap (MAPPIT) assay, in which activation of cytokine receptor activity is used as an readout for protein interaction [71]. With our AFM analysis we found that ∼25% or 19% of FlyBi PPIs met or exceeded the ‘average’ and ‘best’ LIS thresholds, respectively (Figure 6A). More work is needed to understand the extent to which this reflects false positive/negative data in the FlyBi dataset, sub-optimal sensitivity of the LIS approach, or both.

**Figure 6.**
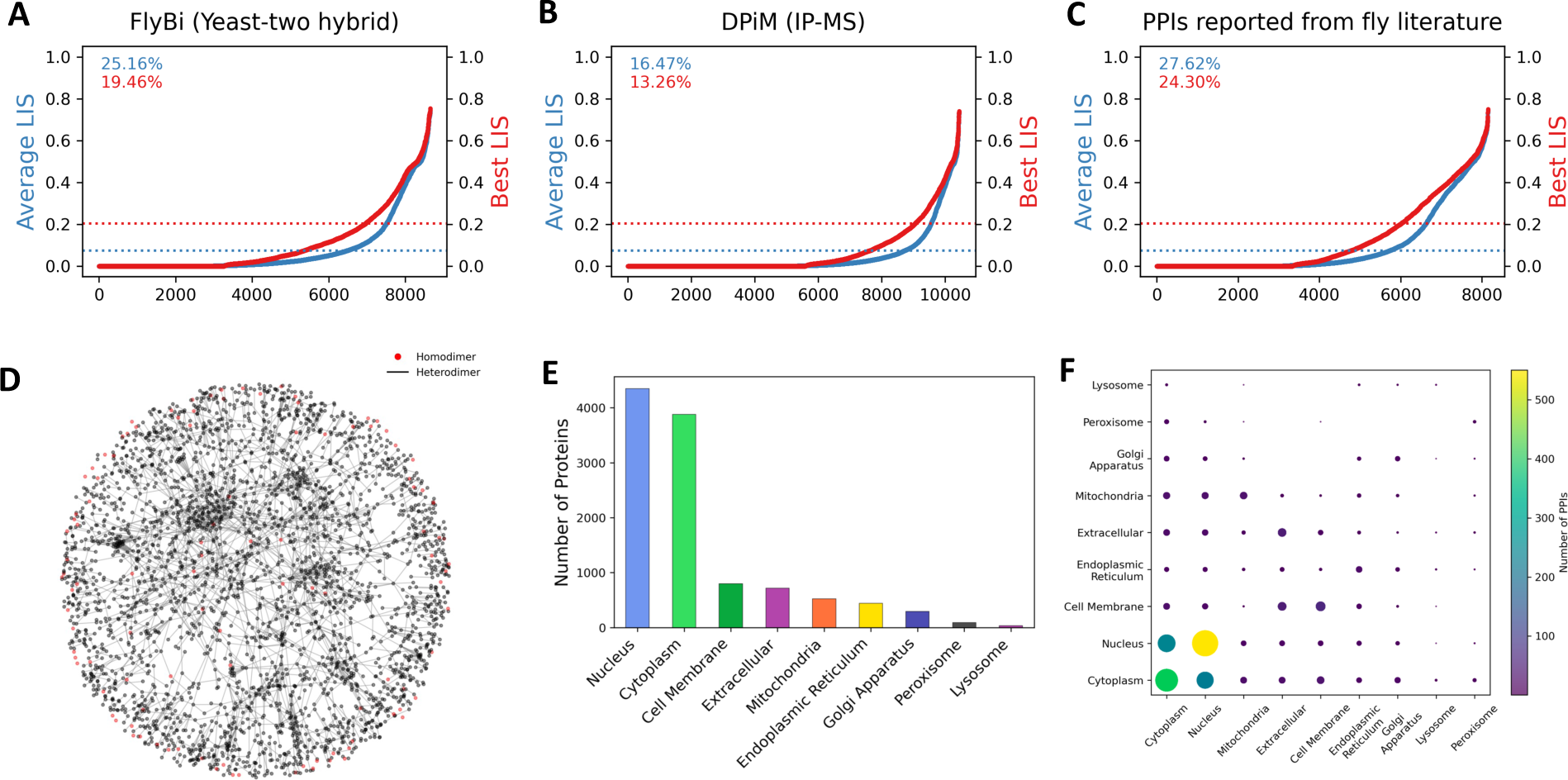
Analysis of *Drosophila* PPI networks using large-scale proteomic data. A. Distribution of the LIS analyzing a comprehensive *Drosophila* Y2H dataset (FlyBi). Percentages at the top of each graph indicate the proportion of PPIs exceeding optimal LIS thresholds indicated by dotted lines. B. Distribution of the LIS analyzing a high-throughput *Drosophila* IP-MS dataset (DPiM). C. Distribution of the LIS analyzing *Drosophila* literature-curated PPIs. D. A network visualization of the *Drosophila* binary interactome from three large datasets, with red nodes representing homodimeric interactions. The network density and the positioning of nodes provide insights into the complexity and connectivity of the proteomic interactions. E. Bar graph depicting the number of proteins categorized by their subcellular localization, as predicted by DeepLoc 2.0 analysis. F. Scatter plot illustrating the distribution of PPIs across different subcellular localizations.

We next analyzed data in the *Drosophila* Protein Interaction Map (DPiM) project, which used IP-MS to identify over 10,000 PPIs using roughly 5,000 bait proteins expressed in *Drosophila* S2R+ cell lines [7]. As expected, the DPiM project identified previously characterized and potential new complexes. Using LIS for predictive analysis within DPiM dataset pairs indicated that ∼ 16% or 13% of the PPIs are predicted to be direct interactions, based on the average and best LIS thresholds, respectively (Figure 6B). The lower percentage of predicted direct interactions in the IP-MS dataset compared to Y2H suggests that PPIs derived from Y2H are more likely to represent direct interactions, aligning with common perception.

Finally, we analyzed a third dataset, comprised of PPIs curated from the *Drosophila* literature (publication dates up to 2021)[11]. This dataset includes >8,000 PPIs selected for inclusion in the dataset because the PPI had been validated using multiple experimental approaches and/or by multiple research groups, such that they were higher confidence PPIs than the set of all PPIs reported in the literature. Subjecting this literature-derived dataset to LIS analysis, we found that 28% and 24% of PPIs met our thresholds for average and best LISs, respectively, that we use as a cutoff for prediction of a direct PPI (Figure 6C). As expected, this rate of recovery surpasses what we found for the large-scale Y2H and IP-MS datasets, underscoring the idea that PPIs corroborated by multiple methods and/or sources are likely to be valid and, specifically, likely to be in spatial proximity to one another, increasing the likelihood that this set of PPIs is enriched for direct interactions.

In our extensive evaluation across three datasets, we uncovered numerous instances of positive PPIs assigned low ipTM scores. Notably, in the FlyBi dataset alone, over 200 interactions were characterized by such low scores, with selected examples detailed in Supplementary Figure 15. A similar trend was observed in the DPiM dataset with over 100 interactions (Supplementary Figure 16), and in the *Drosophila* literature-derived dataset with over 300 interactions (Supplementary Figure 17), suggesting a noteworthy occurrence of PPIs with small interfaces. These findings highlight the power of LIS in capturing nuanced interactions that may be transient or localized yet hold functional significance.

Building upon this insight, we applied the LIS and LIA thresholds to construct network maps of predicted PPIs, offering a comprehensive view of direct interactions (Figure 6D). Prediction of subcellular localization using DeepLoc 2.0 [72] revealed that most proteins in interaction pairs are predicted to be localized to nucleus or in the cytoplasm (Figure 6E). This is in good agreement with the experimental approaches used in the IP-MS and Y2H studies, which we expect are biased towards detection of soluble proteins. Conversely, interactions occurring in the extracellular or membrane environment were less frequent (Figure 6G), consistent with the difficulty inherent to experimental studies of proteins in those regions. Nevertheless, our AFM-based simulations successfully revealed known interactions for some known or predicted membrane-localized and secreted proteins (Supplementary Figure 18), including interactions involving the *Drosophila* Epidermal growth factor receptor (EGFR) and its ligands (Spi, Grk, and Krn) (Supplementary Figure 19). This suggests that computational predictions have the potential to overcome experimental obstacles and identify potential new interactions.

### Development and Expansion of the FlyPredictome Database

To help researchers mine the *Drosophila* PPI predictions described in this work, we have developed FlyPredictome (www.flyrnai.org/tools/fly_predictome), a dedicated online database presented in Figure 7A. This online platform serves as a repository for a wide range of PPI predictions generated through our AFM analysis, offering researchers a valuable tool for investigating PPIs-of-interest.

**Figure 7.**
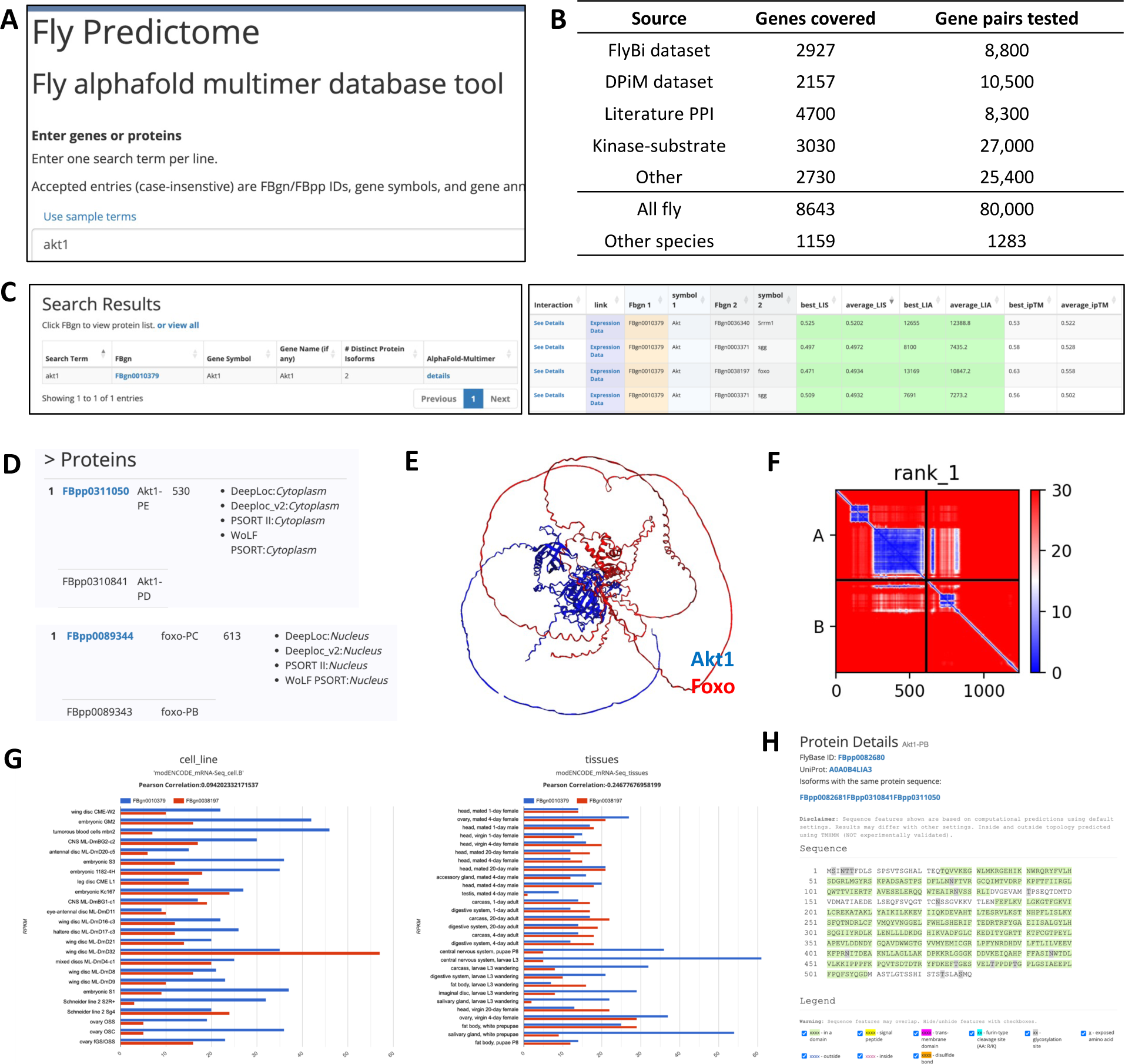
Overview of the FlyPredictome database. A. The main search interface of the FlyPredictome database, where users can enter one or multiple gene or protein names to retrieve PPI data. The database is available at www.flyrnai.org/tools/fly_predictome. B. The summary table of PPI data in FlyPredictome lists various sources of protein-protein interaction (PPI) data, the number of genes each dataset covers, and the number of gene pairs tested. The database features predictions for over 80,000 fly PPIs covering more than 8,600 fly genes, in addition to predictions for over 1,200 PPIs for other species covering more than 1,100 genes. C. Display of search results related to fly Akt1. Prediction results are accessible via the ‘details’ link in AlphaFold-Multimer column, presenting all predictions associated with Akt1. Detailed information for individual predictions is available under the ‘See Details’ link in the Interaction column. If best and average LIS/LIA exceed optimal thresholds, they are highlighted. D. Protein information shows the different isoforms of a protein and their predicted subcellular localizations. E. Protein complex structure is available on the interaction prediction page. F. The PAE map visualizes the confidence of the predicted protein complex structure. G. Graphical representation of expression correlation across various cell lines and tissues. This data can be accessed via the ‘Expression Data’ link on the Search Results page. H. Protein Details page provides comprehensive details for an individual protein, including conserved domains, cleavage sites, transmembrane domains, and post-translational modifications. This detailed view can be accessed by clicking on a specific protein isoform listed in (D).

While a comprehensive proteome-by-proteome prediction is desired, as done in large-scale Y2H experiments, it would be ideal to have a complete coverage for all possible PPIs. However, this is a daunting task as this would require more than a hundred million predictions, which is currently beyond our computational capacity. Thus, we envision a step wise more focused approach. We first prioritized the proteins within the fly major signaling pathways (MSPs) (such as Pvr, Insulin-like, EGFR, FGFP, Sevenless, Torso, Toll, Wnt, BMP, Notch, TNFalpha, Hedgehog, JAK-STAT, Activin, Hippo, and Imd pathways annotated at FlyPhoneDB [73]) as well as key metabolic pathways (as annotated at KEGG). This prioritized strategy has allowed us to expand upon and explore new PPIs within MSPs, including interactions involving receptors, ligands, and other proteins within extracellular or membrane contexts (detailed in Supplementary Figure 20). Moreover, we also analyzed 49 kinases and their potential substrates, with an overview of the results presented in Supplementary Figure 21 and a detailed screening for Akt1 and its substrates shown in Supplementary Figure 22. Currently, the FlyPredictome includes over 80,000 PPI predictions, covering 8,643 *Drosophila* genes, while there are also 1,283 interactions for 1,159 genes for other species (Figure 7B). We plan to expand the database with regular updates on monthly basis.

The FlyPredictome user interface includes a straightforward search function, enabling users to retrieve prediction results for a given protein of interest (Figure 7C-H). At FlyPredictome, users can input a protein and view a list of PPIs predicted using our approach, along with protein-centric and other information such as subcellular location predictions (made using DeepLoc 1.0 [74], DeepLoc 2.0 [72], PSORT II [75], and WoLF PSORT [76]) and co-expression data [77]. In addition, for any given predicted protein pair, users can interact with the predicted structure and PPI interface in a 3D model, as well as highlight specific sub-regions within a model.

Altogether, the information provided should help a user assess if a given predicted PPI is valid and relevant to their study (e.g., co-expressed in a given tissue of interest), design perturbations predicted to disrupt the interaction (e.g., mutations in the predicted interface), and otherwise make use of the predictions to develop and test new hypotheses.

## Discussion

Our study introduces the LIS to improve upon existing AFM metrics such as ipTM [17], pDockQ [24, 53] and pDockQ2 [29]. While these established metrics excel at identifying strong, stable interactions typically found within the cells, they often struggle with the more variable interactions occurring in flexible protein regions, such as IDRs. Departing from the conventional reliance on physical proximity of amino acids, as is the case for pDockQ and pDockQ2, LIS focuses on predictive confidence to enhance the precision of interaction predictions, particularly in flexible regions.

The efficacy of LIS is demonstrated through analysis of positive reference sets from various organisms, ELM-annotated protein pairs, and three extensive datasets (FlyBi, DPiM, fly literature-derived PPIs), revealing its ability to predict interactions between protein domains and flexible regions effectively. Notably, many PPIs with low ipTM but relatively high LIS scores were identified, indicating LIS’s capability in capturing interactions involving flexible regions, including IDRs. Given that a significant portion of protein sequences are predicted to be IDRs and are known to play crucial roles in dynamically regulated interactions [44, 45, 78–81], the ability of LIS to detect such interactions is a significant advancement.

Optimal LIS/LIA thresholds are established through analysis of positive reference sets derived from fly and human literature. However, these reference sets may lean towards stable interactions, potentially limiting the representation of the full spectrum of physiological PPIs.

Challenges arise in cases where the breadth of the interaction area mismatches with the LIS score, leading to potential misclassification of PPIs. Hence, there is a necessity for more customized thresholds to accurately represent the complex reality of PPIs.

Additionally, our analysis reveals potential false positives, where PPIs meeting LIS and LIA thresholds are not supported by predicted subcellular localizations. For example, some PPIs identified in the FlyBi Y2H dataset between proteins from different cellular compartments (e.g., extracellular space vs nucleus) were predicted by LIS analysis to be direct interactions, indicating a potential topological mismatch and underscoring the importance of considering full biological context to avoid incorrectly predicting interactions not occurring in vivo.

Our study establishes the FlyPredictome website, a resource integrating AFM prediction results with data on gene expression and subcellular localization. This comprehensive approach enhances our ability to confirm the biological relevance of predicted PPIs, aiming to reduce false positives and inspire more refined hypotheses for experimental validation.

Lastly, reanalyzing previously reported AFM screens using LIS-based evaluation is anticipated to uncover PPIs missed due to reliance on structural accuracy metrics, thereby enriching our understanding of the PPI landscape and advancing computational strategies in this field.

## Methods

### AlphaFold-Multimer prediction via LocalColabFold

For AFM predictions, we used LocalColabFold (version 1.5.2) [82], incorporating AFM version 2.3.1 with multiple sequence alignments generated through MMseqs2. A variety of Nvidia GPUs (TeslaM40, TeslaV100, RTX6000, RTX8000, A40, A100, and L40) in the Harvard O2 cluster, a high computing environment, were used. Protein complexes of varying sizes were allocated appropriate GPU resources. Consistency in prediction results was observed across different GPUs, with exceptions noted for large protein complexes affected by GPUs with limited memory. The script used for LocalColabFold was as follows:

> Colabfold_batch –num-recycle 5 \
>
> --num-models 5 \
>
> --model-type auto \
>
> --rank multimer \ fasta_file_location \
>
> result_location

Ranks were determined by multimer parameter, ipTM. All ranks were used to calculate average metrics; rank 1, determined by multimer parameter, ipTM, was used to calculate best metrics.

### Calculation of Local Interaction Score and Local Interaction Area

LIS and LIA were calculated using ColabFold output JSON files. Amino acid contacts within the cutoff PAE values were identified as the LIA. Inverted PAE values within the LIA were averaged to determine the LIS. The ‘best’ LIS was derived from the rank 1 model, while ‘average’ LIS was computed from ranks 1-5. A cutoff PAE value of 12, determined to provide the highest AUC, was used for both average and best LIS as per ROC analysis. A Jupyter Notebook for calculating LIS and LIA is available on GitHub (https://github.com/flyark/AFM-LIS).

### Protein-protein interaction reference sets

Curated PPI reference sets for fly, human, and yeast (fly PRS, fly RRS, human PRS, human RRS, yeast PRS, and yeast RRS) [4, 6, 11] were utilized. Random reference sets were treated as negative controls. The longest isoforms for *Drosophila* and reviewed protein sequences for yeast and human datasets were obtained from UniProt. PPIs with unavailable sequences were excluded. Additional negative controls were generated by pairing GFP tagged with two recently reported peptide tags [83], or the fly Wg protein, with proteins from each PRS.

### ELM protein pairs

Protein-linear peptide complexes were identified from the ELM database (http://elm.eu.org/pdbs/). Full-length sequences corresponding to PDB IDs were retrieved from UniProt for AFM prediction. Only pairs yielding two UniProt IDs were processed; those with multiple IDs were excluded.

### Potential kinase-substrate interactions

To make the list of potential kinase-substrate interactions, we used MIST [69] that sources known phosphorylation sites from established database such as PhosphoSitePlus [84] and Phosida [85]. Subsequently, the NetPhorest algorithm [86] was applied to infer interactions based on the linear motifs associated with these phosphorylation sites, excluding low-quality predictions. We focused on 49 kinases and their substrates for AFM prediction. To optimize the computation time, only 101 amino acid sequences from the potential substrates extending 50 amino acids upstream and downstream from the putative phosphorylation sites were used for the analysis.

### Receiver operating characteristic (ROC) analysis

ROC analysis was conducted using the sklearn library in Python, with AUC values computed using positive sets (fly/human PRS) and negative control sets (fly/human RRS, PRS+GFP, PRS+Wg). Optimal thresholds were established based on the highest Youden’s Index. Figures were generated using pandas, NumPy, and Matplotlib in Python.

### Structure visualization

Protein structures were visualized with ChimeraX, using parameters for color-coding by chain, setting the background to white, enabling silhouettes, and applying flat lighting. AlphaFold PAE connections were highlighted using specific JSON files and color domains. The script used for identifying interacting domains was as follows:

> open “pdb file address”
>
> color bychain
>
> set bgColor white
>
> graphics silhouettes true
>
> lighting flat
>
> alphafold pae #1 file “json file address”
>
> alphafold pae #1 connectMaxPae 11
>
> alphafold pae #1 colorDomains true

### Sequential AFM analysis

Multi-stage AFM screening was applied to high-confidence candidates from the IP-MS dataset [47]. LIS and LIA thresholds from ROC analysis were used for stringent filtering, categorizing PPIs as direct or indirect interactions. The process involved sequential screens of baits against their respective preys followed by merging direct PPIs to form comprehensive interactomes.

*The first AFM screen*

> *Bait_1 vs Preys of Bait_1*
>
> *Bait_2 vs Preys of Bait_2*
>
> *Bait_3 vs Preys of Bait_3*
>
> *Bait_4 vs Preys of Bait_4*

*The second AFM screen*

> *Direct interactome* from the first AFM vs Direct interactome* from the first AFM*
>
> *Direct interactome* from the first AFM vs Indirect interactome from the first AFM*

*The third AFM screen*

> *Direct interactome** from the second AFM vs Direct interactome** from the second AFM*
>
> *Direct interactome** from the second AFM vs Indirect interactome from the second AFM*

*: *Direct PPIs from the first AFM screen was merged into direct interactome*

**: *Direct PPIs from the second AFM screen was merged into direct interactome*

### PPI network analysis

Cytoscape was used to visualize PPI networks. Networks in Figure 4D-E were arranged using yFiles tree layouts, and Supplementary Figure 14 was arranged using yFiles radial layouts. Edge thickness indicates average LIS values. The interactome network in Figure 6D was created with Python libraries including pandas, NumPy, networkx, and Matplotlib.

### Construction of the FlyPredictome website

FlyPredictome (https://www.flyrnai.org/tools/fly_predictome/) was built following a three-tier model, with a web-based user-interface at the front end, a database at the backend, and a business logic in the middle tier communicating between the front and back ends by matching input genes with prediction data, expression data, and protein annotations, and retrieving the visualization graphs. The front page is written in PHP using the Symfony framework and front-end HTML pages using the Twig template engine. The JQuery JavaScript library is used to facilitate Ajax calls to the back end, with the DataTables plugin for displaying table views. The Bootstrap framework and some custom CSS are used on the user interface. 3Dmol.js is used for 3D structure visualization of the evaluated protein pairs [87]. A mySQL database is used to store the prediction results, transcriptomic data as well as the knowledgebase of protein annotation. Both the website and databases are hosted on the O2 high-performance computing cluster, which is made available by the Research Computing group at Harvard Medical School.

The knowledgebase of protein information was built by retrieving information from various public resources or by doing the prediction using published tools. Specifically, the protein sequences were retrieved from FlyBase and RefSeq while protein annotations such as protein domain and post-translational modification were retrieved from UniProt, netCglyc, netNglyc, and netOglyc of the glycolysylation sites [88, 89]. Sub-cellular localization predictions were done using PSORT II [75], WoLF PSORT [76], DeepLoc 1.0 [74], and DeepLoc 2.0 [72]; ProP 1.0 was used for predicting signal peptide and Furin-cleavage sites [90]; and TMHMM 2.0 was used for predicting transmembrane helices in proteins [91].

## Supporting information

Supplementary Table related to Figure 3

Supplementary Table_Positive PPIs

Supplementary Table_total PPI prediction

## Acknowledgements

We are grateful to the Research Computing Group at Harvard Medical School for access to the O2 High Performance Compute Cluster. We thank Weihang Chen at DRSC for helping with DeepLoc v2 prediction and Bernard Mathey-Prevot and Joshua Shing Shun Li for commenting on the manuscript. This research was supported by NIH NIGMS P41 GM132087 and NIH NIAMS R01 AR057352. AK was supported by Postdoctoral Fellowship Program (Nurturing Next-generation Researchers) through the National Research Foundation of Korea (NRF) funded by the Ministry of Education (2021R1A6A3A14039622). NP is an investigator of Howard Hughes Medical Institute.

This article is subject to HHMI’s Open Access to Publications policy. HHMI lab heads have previously granted a nonexclusive CC BY 4.0 license to the public and a sublicensable license to HHMI in their research articles. Pursuant to those licenses, the author-accepted manuscript of this article can be made freely available under a CC BY 4.0 license immediately upon publication.

## Declaration of interests

The authors declare no competing interests.

## Declaration of generative AI used in the writing process

During the preparation of this manuscript, the authors used OpenAI’s ChatGPT to assist in generating custom Python scripts for the analysis of prediction results and to improve the readability and language. After using this tool, the authors reviewed and edited the content as needed and take full responsibility of the publication.

## Notes

https://www.flyrnai.org/tools/fly_predictome

### Competing Interest Statement

The authors have declared no competing interest.

