## Supplementary Figure 1-22 for "Enhanced Protein-Protein Interaction Discovery via AlphaFold-Multimer"

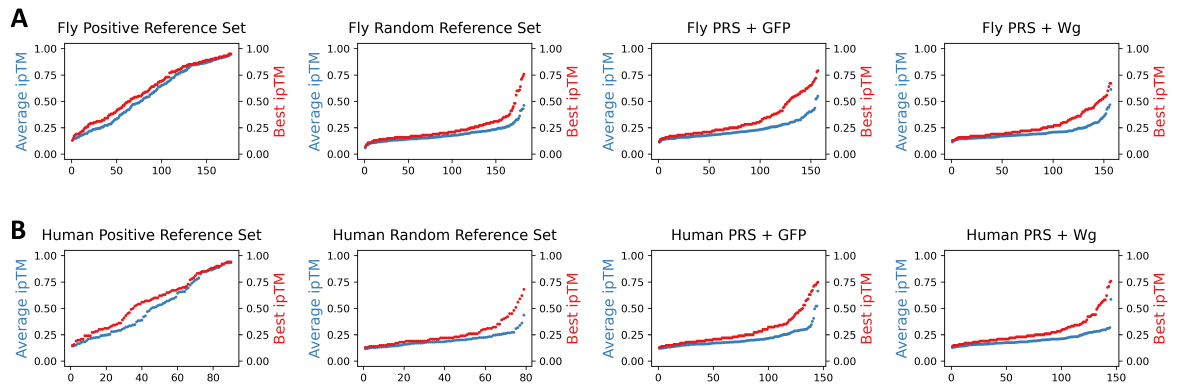

**Supplementary Figure 1. Distribution of average and best ipTM scores for fly and human positive and control PPI sets.**

A. Distribution of average and best ipTM scores for fly positive PPIs (Fly PRS) and fly control PPIs (Fly RRS, Fly PRS + GFP, Fly PRS + Wg).
B. Distribution of average and best ipTM scores for human positive PPIs (human PRS) and human control PPIs (human RRS, human PRS + GFP, human PRS + Wg).

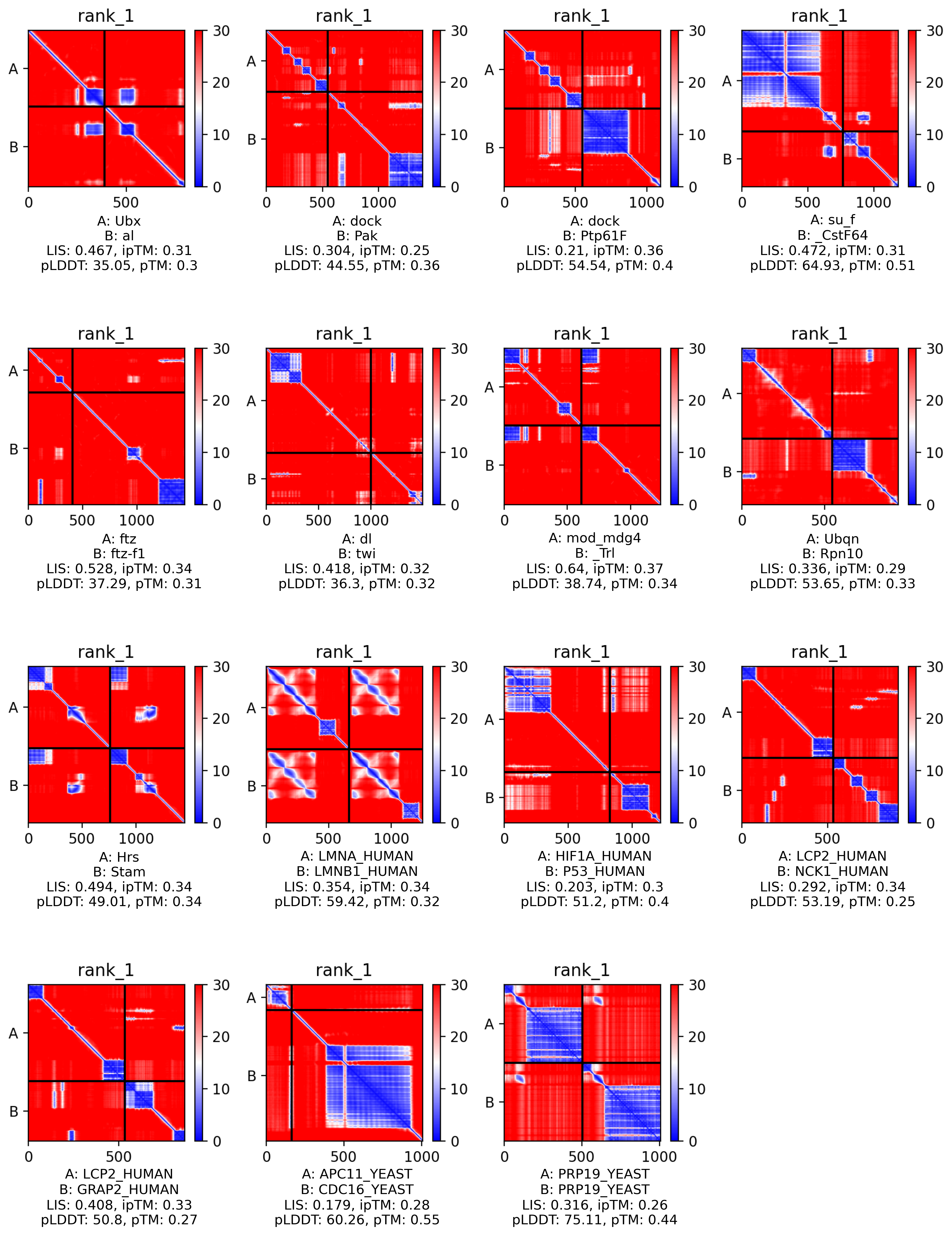

**Supplementary Figure 2. Example PPIs with low ipTM Scores but distinct PAE map interactions.**

Examples of PPIs with low ipTM scores display pronounced blue areas on PAE maps. Despite the overall low scores for pLDDT and pTM, indicative of structural uncertainty, these PPIs exhibit clear interaction zones. Notably, the interactions appear to involve either two well-defined rigid domains or a rigid domain interacting with a flexible region across both protein partners. Additionally, each protein in these pairs typically features extended flexible regions.

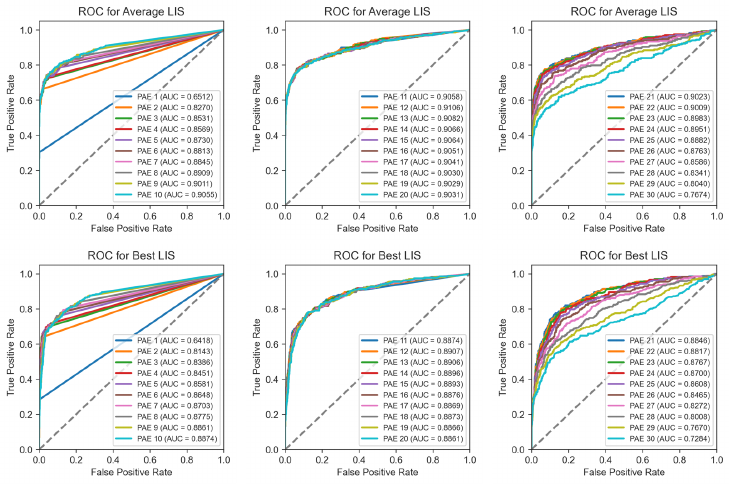

**Supplementary Figure 3. ROC analysis for determining optimal PAE cutoff in LIS calculation.**

This series of Receiver Operating Characteristic (ROC) curves evaluates the LIS across a range of PAE cutoff values. The analysis assesses both average and best LIS metrics to establish the PAE cutoff that maximizes the Area Under the Curve (AUC), indicating optimal balance between sensitivity and specificity in PPI prediction. Positive PPI sets, consisting of fly and human Positive Reference Sets (PRSs), served as true positives, while negative sets—comprising fly and human Random Reference Sets (RRSs), PRSs paired with GFP, and PRSs paired with the fly Wingless (Wg) protein—served as false positives. The highest AUC for both average and best LIS metrics was achieved at a PAE cutoff of 12.

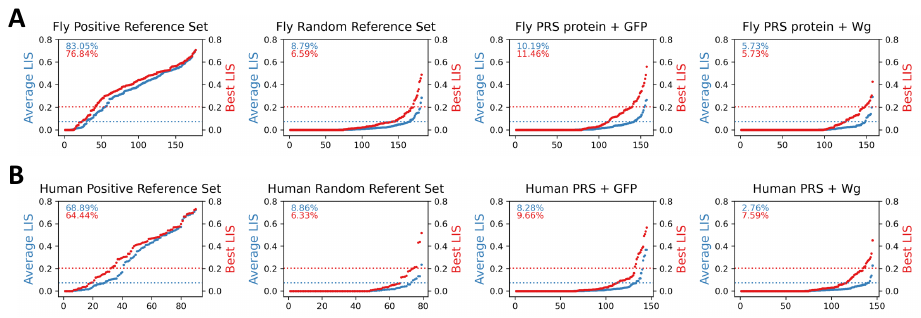

**Supplementary Figure 4. Distribution of average and best LIS values for fly and human positive and control PPI sets**

A. Distribution of average and best LIS values for fly positive PPIs (Fly PRS) and fly control PPIs (Fly RRS, Fly PRS + GFP, Fly PRS + Wg). Percentages at the top of each graph indicate the proportion of PPIs exceeding an optimal threshold (A-B).
B. Distribution of average and best LIS values for human positive PPIs (human PRS) and human control PPIs (human RRS, human PRS + GFP, human PRS + W

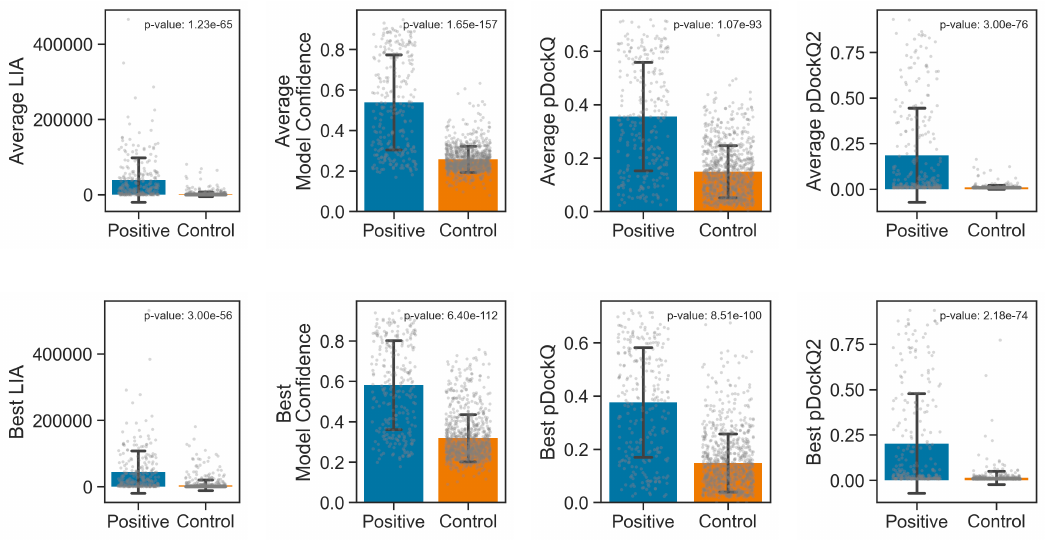

**Supplementary Figure 5. Statistical analysis of average and best metrics for PPI prediction.**

Comparative analysis of the mean scores for both average and best metrics across positive and control PPI sets. The metrics evaluated include LIA, Model Confidence, pDockQ, and pDockQ2. Boxplots represent the distribution of scores within the positive and control PPI datasets. Above each boxplot, the p-values derived from t-tests are displayed, indicating the statistical significance of the differences between the positive and control groups for each metric.

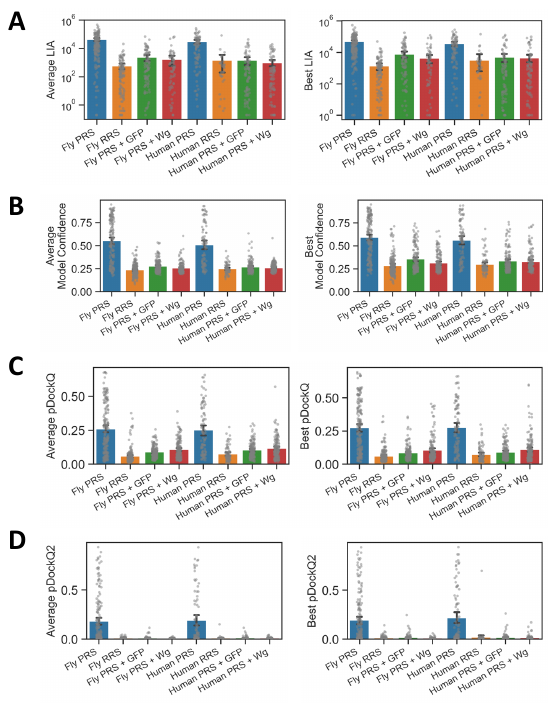

**Supplementary Figure 6. Detailed metric evaluations for individual protein-protein interaction sets.**

A. Boxplots depict the mean average and best LIA scores across various PPI sets, including fly and human Positive Reference Sets (PRS), Random Reference Sets (RRS), and PRS paired with GFP (PRS + GFP) and fly Wingless (Wg) proteins (PRS + Wg)**.**
B. Boxplots illustrate the mean average and best Model Confidence values for the same PPI sets**.**
C. Boxplots show the mean average and best pDockQ scores**.**
D. Boxplots present the mean average and best pDockQ2 scores**.**

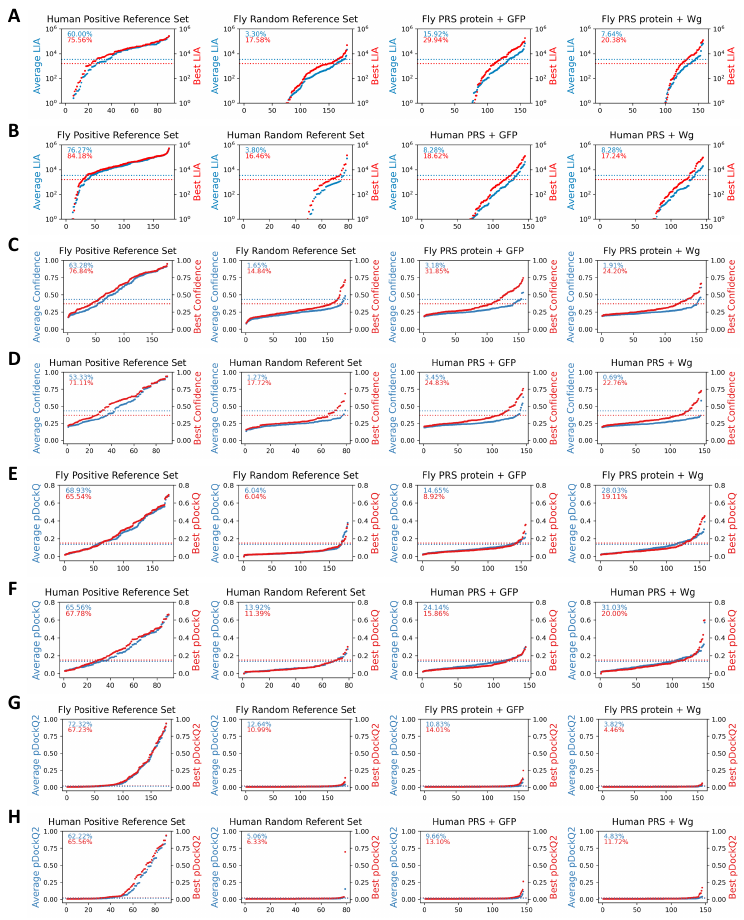

**Supplementary Figure 7. Distribution of each AFM metric for individual PPI sets.**

A. Distribution of average and best LIA across individual fly PPI sets**.** Percentages at the top of each graph indicate the proportion of PPIs exceeding each optimal threshold (A-H)
B. Distribution of average and best LIA across individual human PPI sets**.**
C. Distribution of average and best model confidence across individual fly PPI sets**.**
D. Distribution of average and best model confidence across individual human PPI sets**.**
E. Distribution of average and best pDockQ across individual fly PPI sets**.**
F. Distribution of average and best pDockQ across individual human PPI sets**.**
G. Distribution of average and best pDockQ2 across individual fly PPI sets**.**
H. Distribution of average and best pDockQ2 across individual human PPI sets**.**

| **Metric** | **Optimal Threshold** | **Specificity** | **Sensitivity** | **AUC** | **Youden's Index** |
| --- | --- | --- | --- | --- | --- |
| Average LIS | 0.073 | 0.926 | 0.787 | 0.911 | 0.713 |
| Best LIS | 0.203 | 0.919 | 0.730 | 0.891 | 0.649 |
| Average LIA | 1610.4 | 0.876 | 0.768 | 0.889 | 0.644 |
| Best LIA | 3432 | 0.855 | 0.775 | 0.866 | 0.631 |
| Average ipTM | 0.322 | 0.938 | 0.712 | 0.891 | 0.649 |
| Best ipTM | 0.38 | 0.823 | 0.734 | 0.863 | 0.557 |
| Average Confidence | 0.367 | 0.951 | 0.674 | 0.859 | 0.626 |
| Best Confidence | 0.432 | 0.854 | 0.685 | 0.841 | 0.540 |
| Average pDockQ | 0.133 | 0.805 | 0.682 | 0.796 | 0.486 |
| Best pDockQ | 0.149 | 0.866 | 0.667 | 0.818 | 0.533 |
| Average pDockQ2 | 0.015 | 0.918 | 0.693 | 0.842 | 0.611 |
| Best pDockQ2 | 0.021 | 0.896 | 0.670 | 0.832 | 0.566 |

**Supplementary Figure 8. Assessment of AlphaFold-Multimer metrics using ROC analysis.**

The table summarizes the results from ROC analysis for various AFM metrics. For each metric, the optimal threshold was determined by the highest Youden's Index value, which optimizes the trade-off between true positive and false positive rates. Metrics are evaluated on specificity, sensitivity, area under curve (AUC), and Youden's Index. The average and best values of LIS, LIA, ipTM and model confidence, in addition to pDockQ and pDockQ2, are presented.

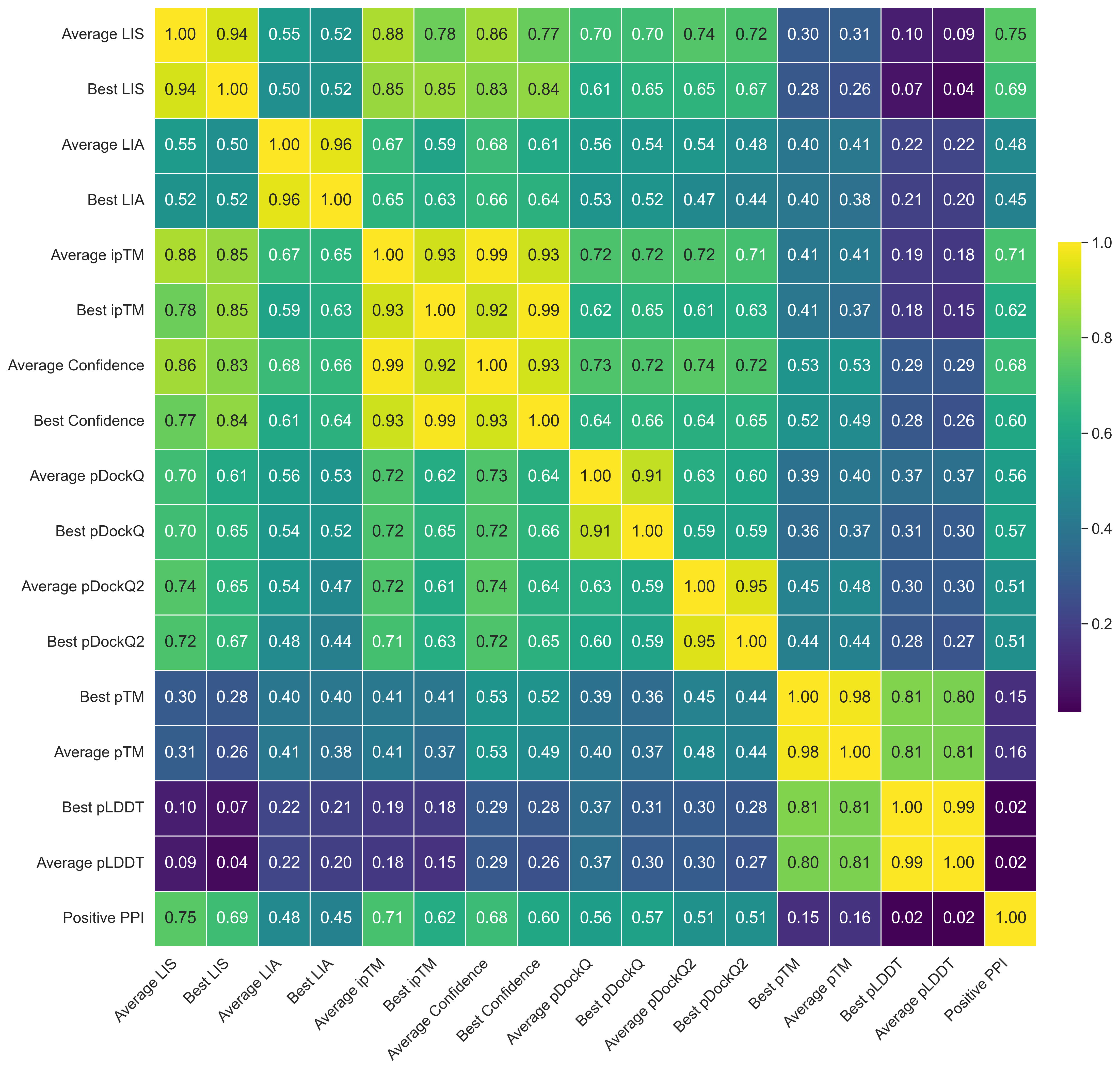

**Supplementary Figure 9. Correlation heatmap for AFM-derived metrics in PPI prediction.**

This heatmap represents the correlation coefficients between various AFM metrics used for PPI prediction. Each cell in the heatmap shows the Pearson correlation coefficient between pairs of metrics, with values closer to 1.0 indicating a stronger positive correlation and those closer to 0 indicating no correlation. Fly and human positive reference sets are indicated as Positive PPI.

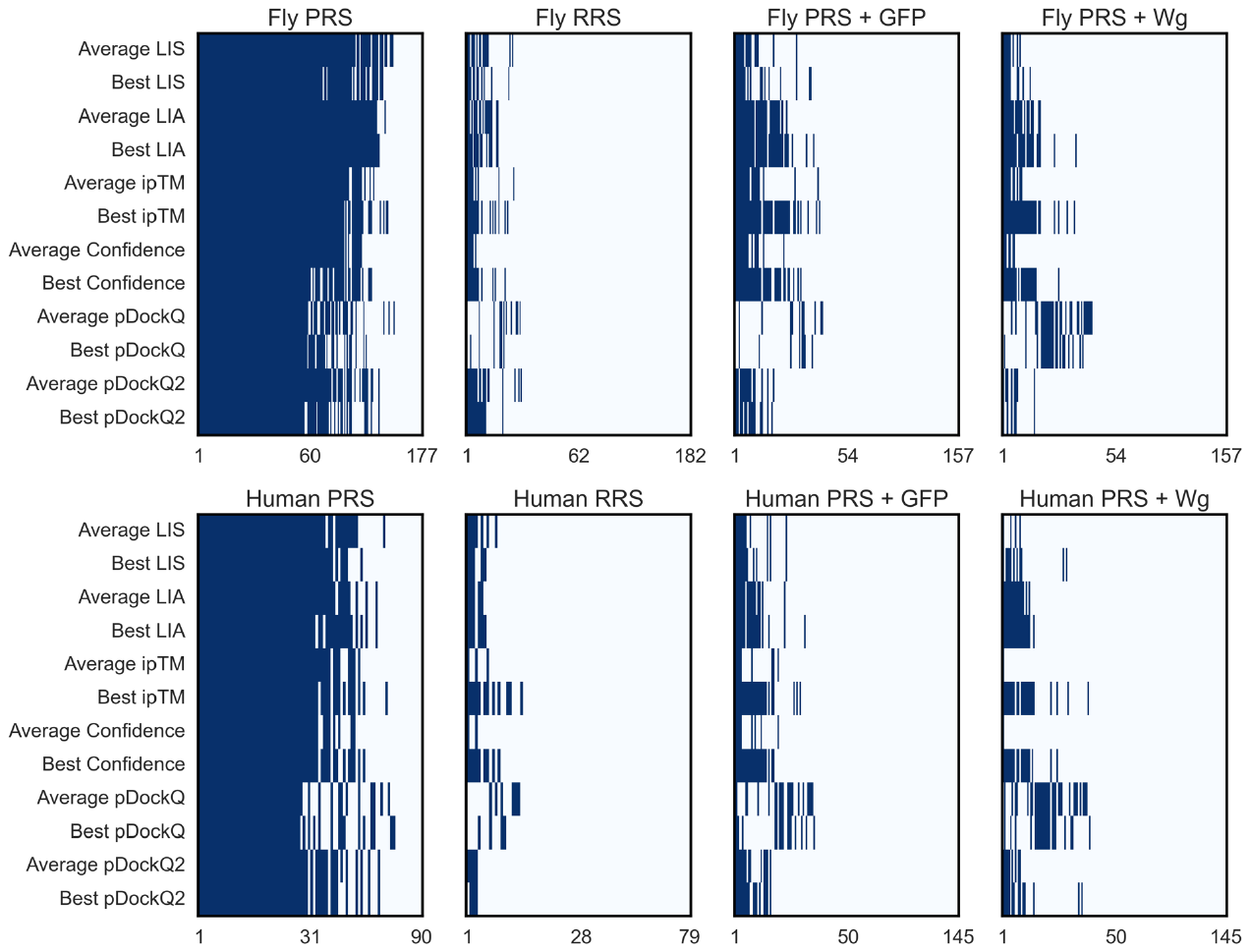

**Supplementary Figure 10. Heatmap of PPI predictions across metrics.**

This heatmap visualizes the predictive power of different metrics by highlighting PPIs that exceed the optimal thresholds established by ROC analysis. The fly and human PPI sets include Positive Reference Sets (PRS), Random Reference Sets (RRS), and PRS paired with GFP (PRS + GFP) or fly Wingless (Wg) (PRS + Wg) as controls. Interactions that surpass the optimal metric thresholds are marked in blue, indicating a predicted direct interaction according to the respective metric.

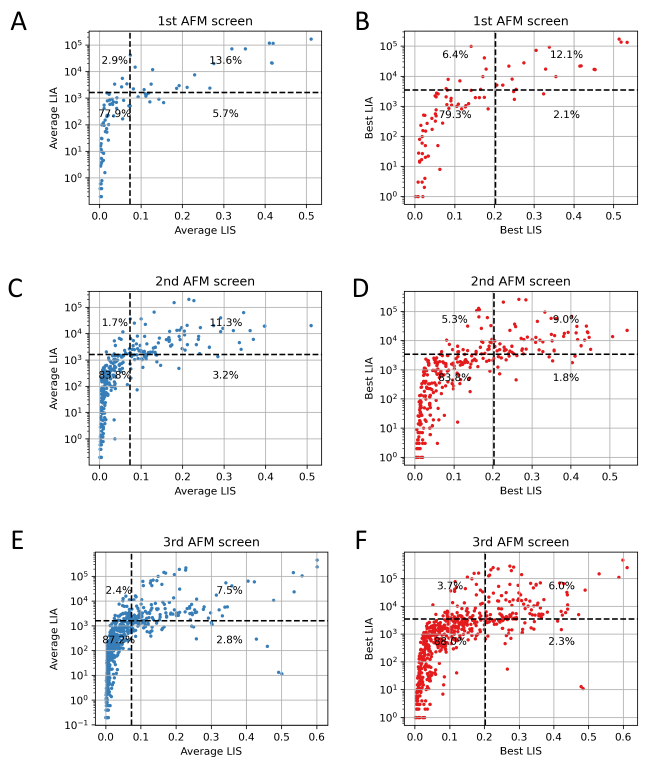

**Supplementary Figure 11. Analysis of m6A interactome via sequential AFM screens.**

These plots present the results from sequential AFM screening rounds applied to an m6A IP-MS interactome dataset. For each round, the average and best LIS are plotted against LIA, with PPIs meeting the optimal thresholds for both metrics predicted as direct interactions.

A. Results from the 1st AFM screen showing the relationship between average LIS and LIA. The optimal thresholds for prediction are marked by black dotted lines, with PPIs above these thresholds highlighted to indicate direct interactions (A-F).
B. Results from the 1st AFM screen for best LIS and LIA.
C. Results from the 2nd AFM screen for average LIS and LIA.
D. Results from the 2nd AFM screen for best LIS and LIA.
E. Results from the 3rd AFM screen for best LIS and LIA.
F. Results from the 3rd AFM screen for average LIS and LIA.

| **Kap-alpha3- interacting proteins** | **Best LIS** | **Average LIS** | **Best ipTM** | **Average ipTM** | **Fly PubMed** | **Ortholog PubMed** |
| --- | --- | --- | --- | --- | --- | --- |
| B52 | 0.544 | 0.5094 | 0.47 | 0.326 |  |  |
| Bacc | 0.507 | 0.3224 | 0.7 | 0.68 |  |  |
| CCT7 | 0.451 | 0.2758 | 0.47 | 0.316 |  |  |
| CG10417 | 0.446 | 0.2386 | 0.73 | 0.672 |  |  |
| CG2199 | 0.443 | 0.3622 | 0.71 | 0.652 | 14605208 |  |
| D1 | 0.417 | 0.1938 | 0.34 | 0.292 |  |  |
| Df31 | 0.413 | 0.3982 | 0.75 | 0.728 |  |  |
| Droj2 | 0.383 | 0.2966 | 0.59 | 0.456 |  |  |
| Fkbp39 | 0.362 | 0.2916 | 0.75 | 0.622 |  |  |
| Flacc | 0.358 | 0.2912 | 0.72 | 0.584 |  |  |
| Hrb98DE | 0.35 | 0.2042 | 0.56 | 0.27 |  |  |
| Lam | 0.328 | 0.1412 | 0.7 | 0.63 |  |  |
| METTL14 | 0.32 | 0.1652 | 0.45 | 0.342 |  |  |
| METTL3 | 0.314 | 0.2744 | 0.72 | 0.652 |  | 26344197, 14704431, 24623722, 29568061 |
| Nap1 | 0.3 | 0.0834 | 0.69 | 0.494 |  |  |
| Nito | 0.274 | 0.1344 | 0.54 | 0.458 |  |  |
| Rm62 | 0.272 | 0.144 | 0.49 | 0.352 |  | 26344197 |
| RpL22 | 0.248 | 0.054 | 0.66 | 0.578 |  |  |
| RpLP0 | 0.233 | 0.081 | 0.62 | 0.416 |  |  |
| Trxr-1 | 0.205 | 0.1684 | 0.65 | 0.354 |  |  |
| fl(2)d | 0.196 | 0.168 | 0.71 | 0.674 |  |  |
| lark | 0.179 | 0.1292 | 0.35 | 0.308 |  |  |
| lost | 0.171 | 0.078 | 0.58 | 0.492 |  |  |
| nudE | 0.168 | 0.0716 | 0.55 | 0.36 |  |  |
| pasha | 0.156 | 0.0896 | 0.74 | 0.576 |  |  |
| rump | 0.122 | 0.0746 | 0.51 | 0.328 |  |  |
| sqd | 0.11 | 0.0706 | 0.64 | 0.454 | 22036573 | 26344197 |

**Supplementary Figure 12. Kap-alpha3-interacting proteins predicted by sequential AFM screening.**

This table lists the proteins predicted to interact with Kap-alpha3 through sequential AFM screening. The 'Fly PubMed' column lists the PubMed IDs of studies that have previously reported these PPIs involving *Drosophila* proteins. The 'Ortholog PubMed' column provides PubMed IDs for studies reporting interactions of orthologous proteins in other species. PubMed IDs were sourced from the MIST database.

| **14-3-3epsilon-interacting proteins** | **Best LIS** | **Average LIS** | **Best ipTM** | **Average ipTM** | **Fly Pubmed** | **Orthlog Pubmed** |
| --- | --- | --- | --- | --- | --- | --- |
| 14-3-3epsilon | 0.587 | 0.5584 | 0.86 | 0.844 | 12431373 |  |
| D1 | 0.434 | 0.3454 | 0.74 | 0.642 |  |  |
| qkr58E-1 | 0.425 | 0.239 | 0.75 | 0.564 |  | 16615898 |
| Calr | 0.41 | 0.341 | 0.74 | 0.674 | 22036573 | 26344197, 16615898 |
| Bacc | 0.371 | 0.2584 | 0.74 | 0.518 |  |  |
| Nap1 | 0.364 | 0.2982 | 0.67 | 0.616 |  |  |
| Df31 | 0.361 | 0.2798 | 0.72 | 0.538 | 22036573 |  |
| qkr58E-2 | 0.361 | 0.322 | 0.7 | 0.668 |  | 16615898 |
| Lam | 0.357 | 0.2332 | 0.73 | 0.634 | 20818332 | 16615898 |
| sqd | 0.357 | 0.171 | 0.71 | 0.41 |  | 16615898, 16944949 |
| CG10417 | 0.346 | 0.191 | 0.72 | 0.568 |  |  |
| CG7903 | 0.331 | 0.1546 | 0.65 | 0.388 |  |  |
| RpLP0 | 0.328 | 0.135 | 0.62 | 0.39 |  |  |
| nudE | 0.318 | 0.1222 | 0.63 | 0.34 | 23987511 | 12796778, 25332407, 16944949, 18331715 |
| Pep | 0.316 | 0.0632 | 0.7 | 0.292 |  |  |
| yps | 0.292 | 0.2586 | 0.61 | 0.506 |  | 16944949, 16615898 |
| RanBPM | 0.285 | 0.0984 | 0.69 | 0.404 |  | 25172955 |
| Hakai | 0.279 | 0.238 | 0.7 | 0.614 |  | 16615898 |
| CG9641 | 0.273 | 0.0938 | 0.55 | 0.372 |  |  |
| fl(2)d | 0.266 | 0.2062 | 0.57 | 0.524 |  |  |
| RpL22 | 0.256 | 0.098 | 0.55 | 0.268 |  |  |
| bel | 0.252 | 0.0518 | 0.65 | 0.298 |  | 16615898 |
| Rm62 | 0.249 | 0.1104 | 0.68 | 0.424 |  |  |
| smt3 | 0.249 | 0.1806 | 0.61 | 0.436 | 26290570 | 16615898 |
| lark | 0.243 | 0.1738 | 0.42 | 0.326 |  |  |
| Fkbp39 | 0.229 | 0.0862 | 0.59 | 0.346 |  |  |
| Flacc | 0.229 | 0.1074 | 0.65 | 0.464 |  |  |
| lost | 0.229 | 0.152 | 0.57 | 0.518 |  |  |
| pAbp | 0.221 | 0.0928 | 0.64 | 0.348 |  | 14744259, 16615898 |
| B52 | 0.216 | 0.2102 | 0.33 | 0.288 |  |  |
| CG7065 | 0.202 | 0.0994 | 0.61 | 0.45 |  |  |
| CCT7 | 0.2 | 0.112 | 0.51 | 0.346 |  | 16615898 |
| CG7611 | 0.177 | 0.1298 | 0.6 | 0.486 |  |  |
| Nito | 0.171 | 0.1338 | 0.53 | 0.412 |  |  |
| sesB | 0.158 | 0.1248 | 0.44 | 0.3 |  |  |

**Supplementary Figure 13. 14-3-3 epsilon-interacting proteins predicted by sequential AFM screening.**

This table lists the proteins predicted to interact with 14-3-3 epsilon through sequential AFM screening. The 'Fly PubMed' column lists the PubMed IDs of studies that have previously reported these PPIs involving *Drosophila* proteins. The 'Ortholog PubMed' column provides PubMed IDs for studies reporting interactions of orthologous proteins in other species. PubMed IDs were sourced from the MIST database.

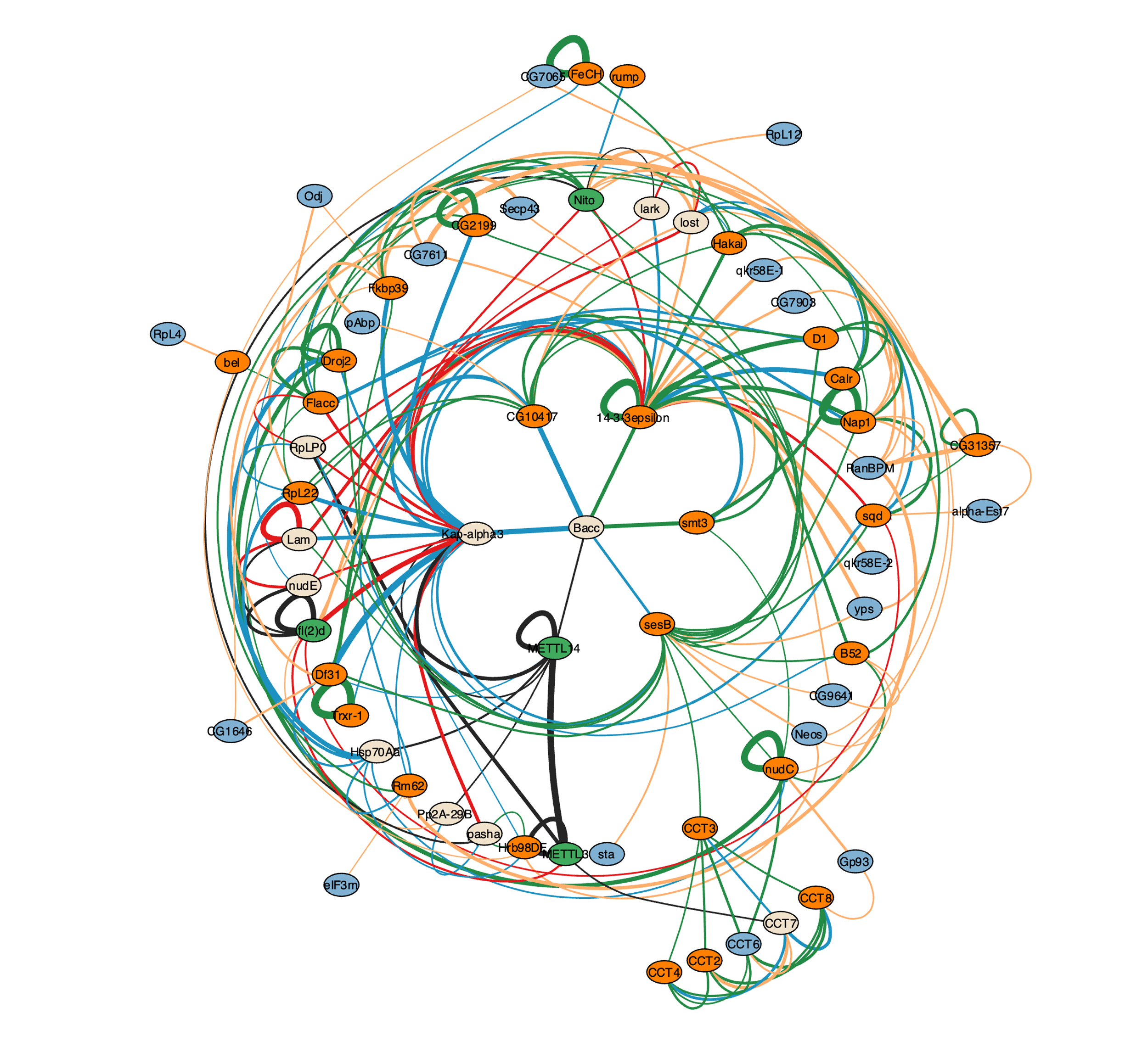

**Supplementary Figure 14. Comprehensive network from sequential AFM screening for *Drosophila* m6A complex IP-MS interactome.**

This comprehensive network map visualizes the direct PPIs within the IP-MS dataset of the *Drosophila* m6A RNA methylation complex, as identified through iterative AFM screening processes. Each node represents a protein, color-coded to indicate its group based on the screening round: primary baits in green, Group 1 proteins in beige, Group 2 in orange, and Group 3 in blue. The edges connecting the nodes represent identified PPIs, with varying thickness corresponding to the average LIS; thicker edges denote higher LIS values, indicative of stronger interaction predictions.

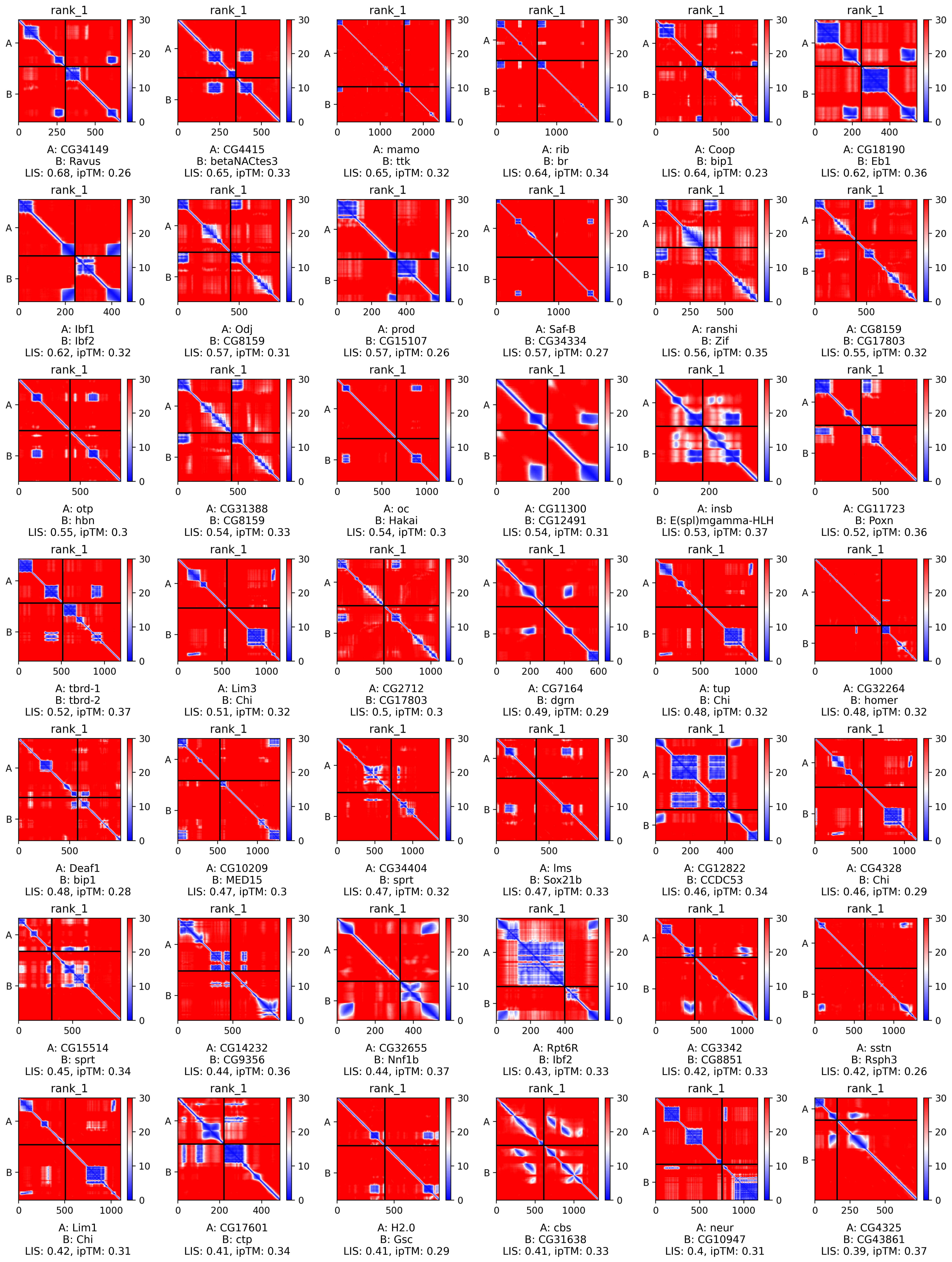

**Supplementary Figure 15. Example positive PPIs with low ipTM scores in FlyBi dataset.**

PAE maps displaying examples of PPIs with small interaction interfaces from FlyBi Y2H dataset. Names of individual proteins, LIS, and ipTM scores are specified for each example.

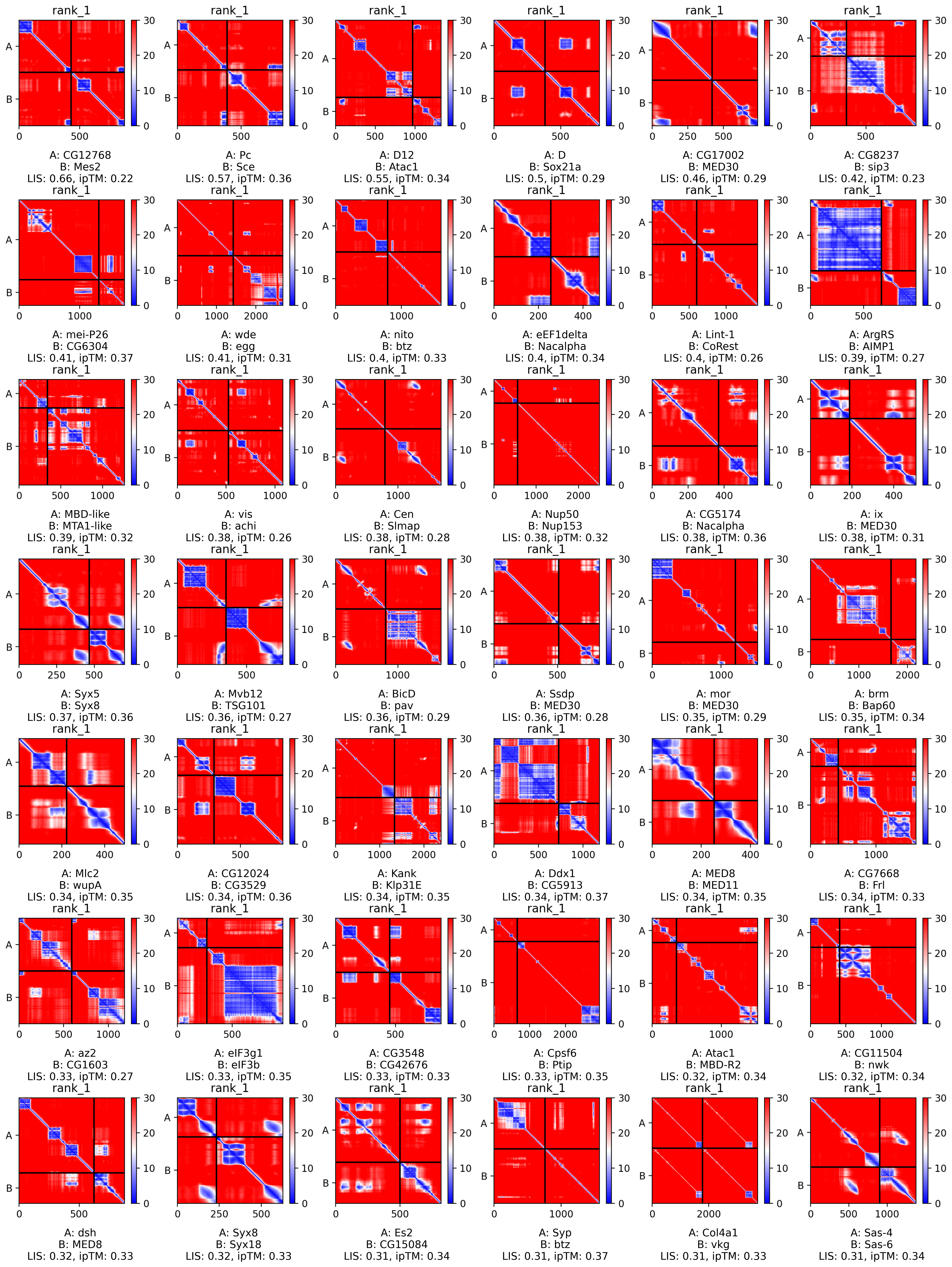

**Supplementary Figure 16. Example positive PPIs with low ipTM scores in DPiM dataset.**

PAE maps displaying examples of PPIs with small interaction interfaces from DPiM IP-MS dataset. Names of individual proteins, LIS, and ipTM scores are specified for each example.

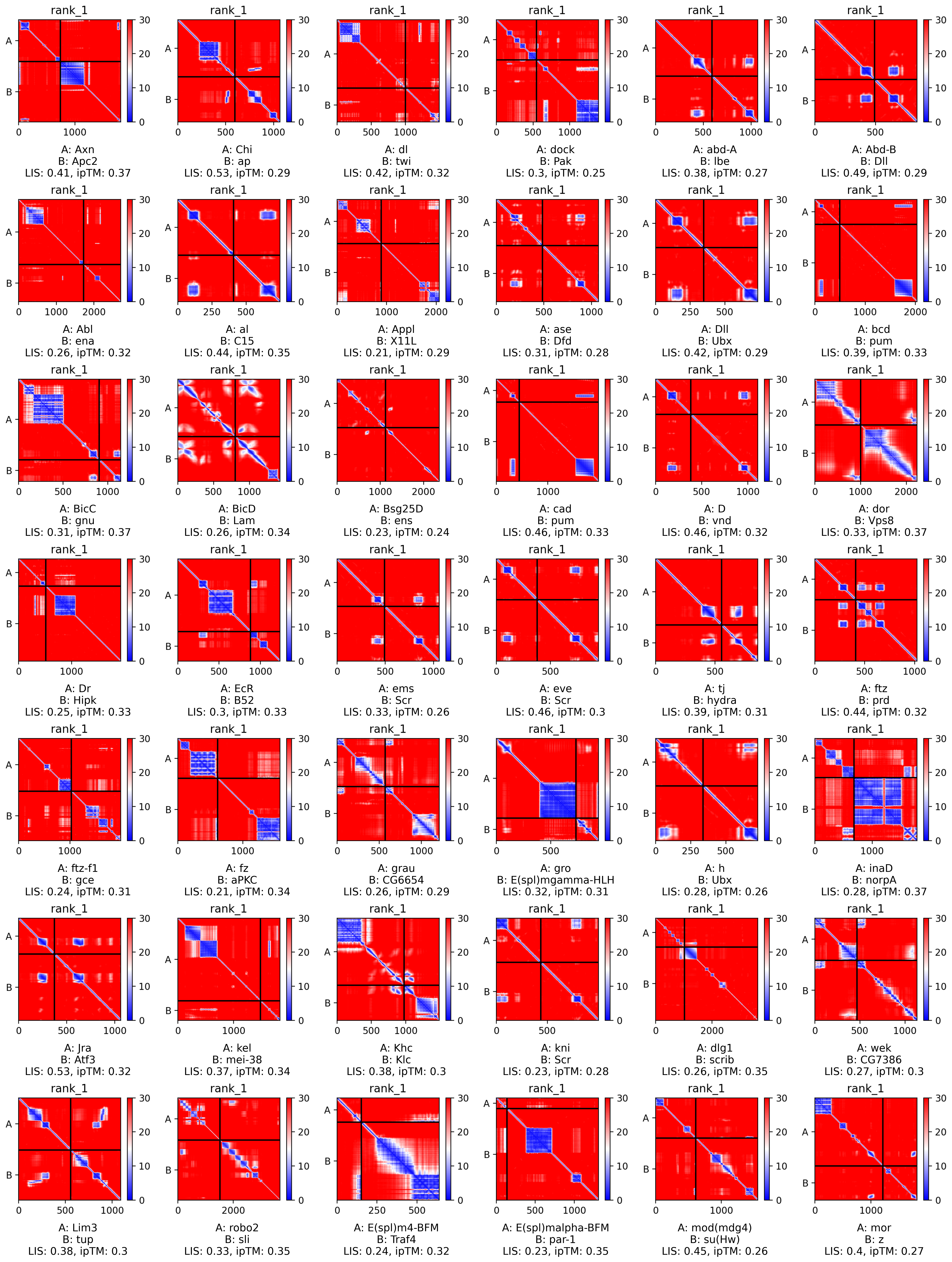

**Supplementary Figure 17. Example positive PPIs with low ipTM scores in fly literature-derived PPIs.**

PAE maps displaying examples of PPIs with small interaction interfaces from fly literature-derived dataset. Names of individual proteins, LIS, and ipTM scores are specified for each example.

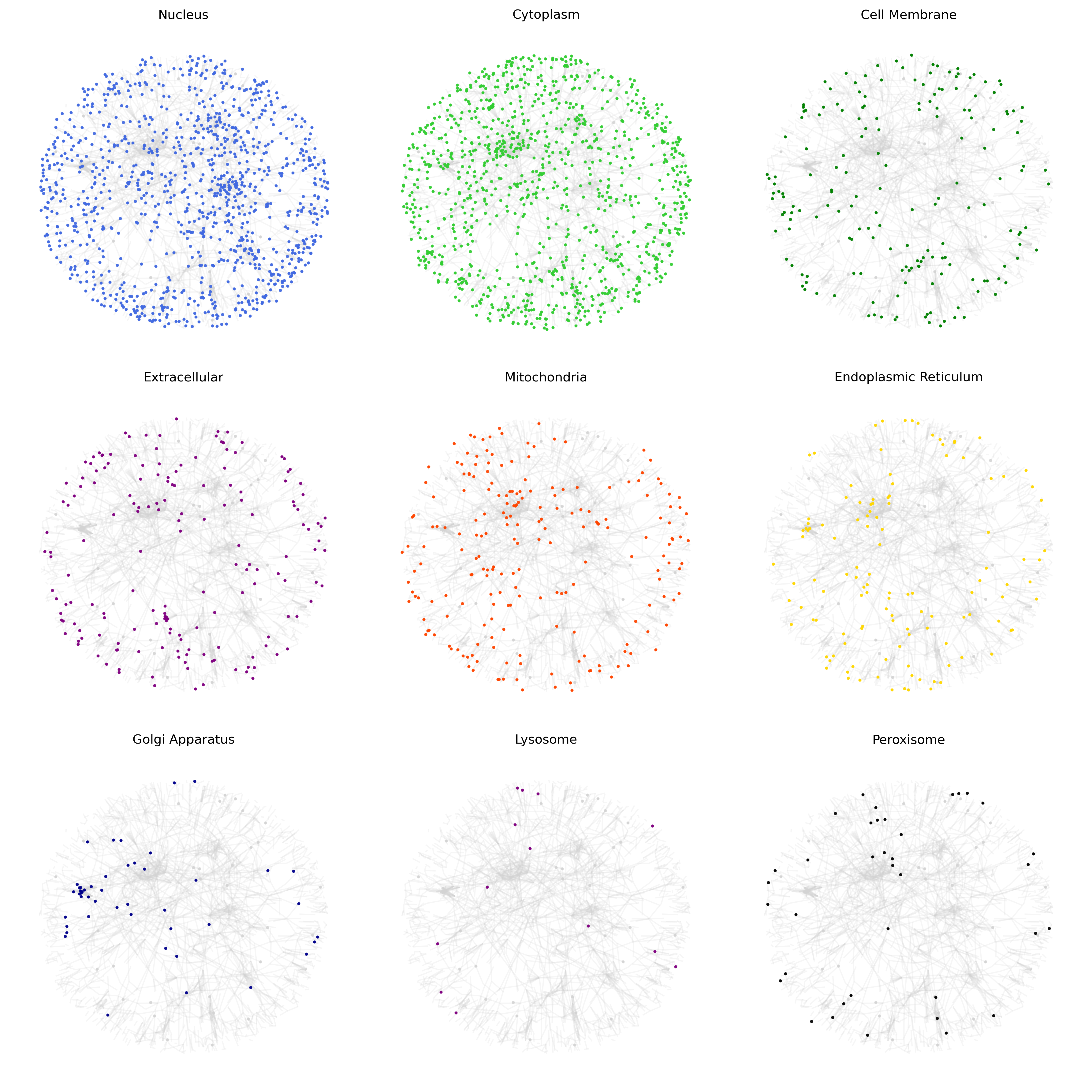

**Supplementary Figure 18. Subcellular Distribution Networks of *Drosophila* Direct PPI interactome.**

Subcellular localization predicted by DeepLoc 2.0 is highlighted in the *Drosophila* PPI network built from three large PPI datasets.

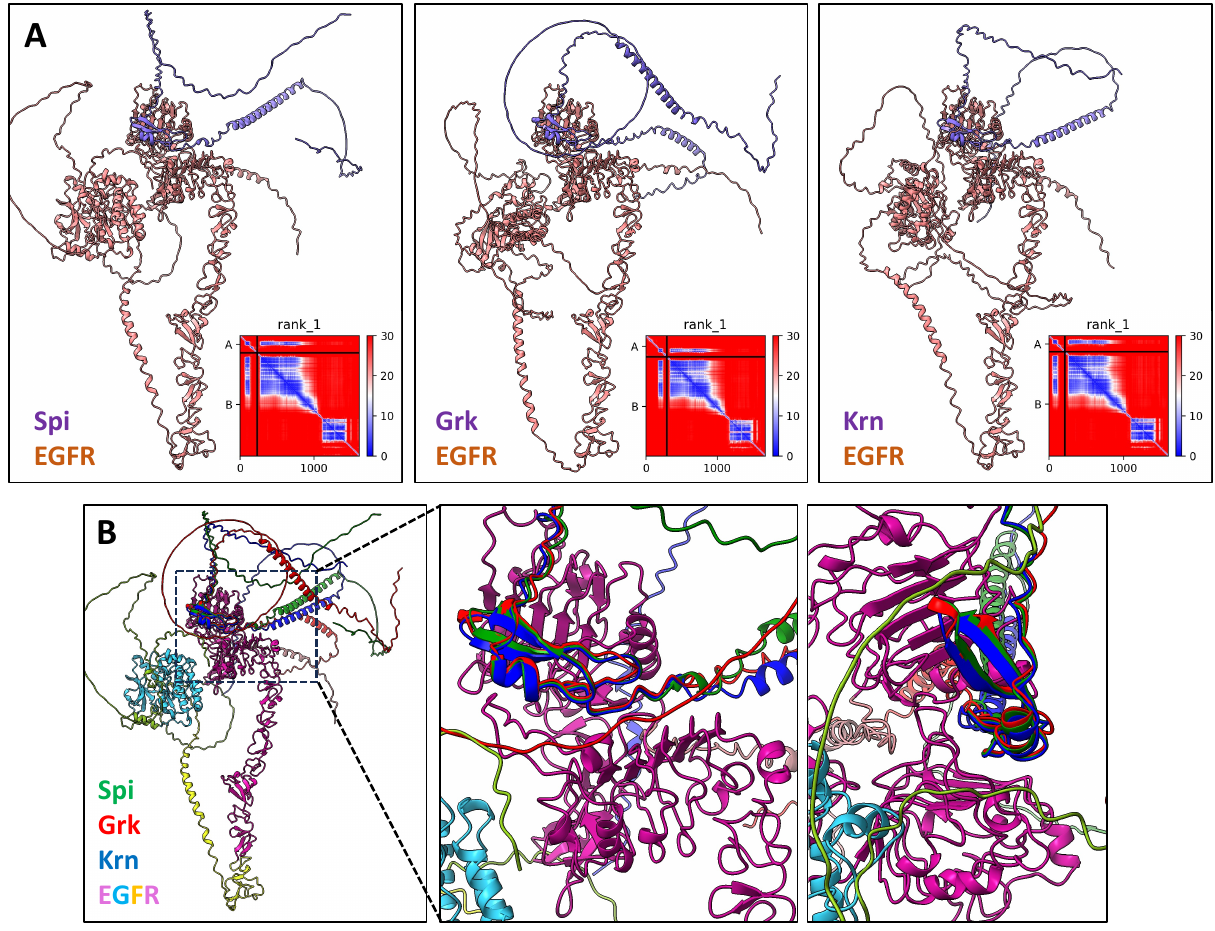

**Supplementary Figure 19. Protein complex structures of *Drosophila* EGFR and EGF ligands (Spi, Grk, and Krn) predicted by AFM.**

A. Structural predictions of *Drosophila* EGFR with EGF ligands Spi, Grk, and Krn using AFM. The EGF ligands are depicted in purple, illustrating their interaction with the receptor, color in pink. PAE map for each prediction is present in the right bottom corner.
B. Detailed view of the ligand-binding interface on EGFR. Spi, Grk, and Krn interact with a conserved binding interface on the EGFR, as indicated by similar spatial arrangements within the predictive models.

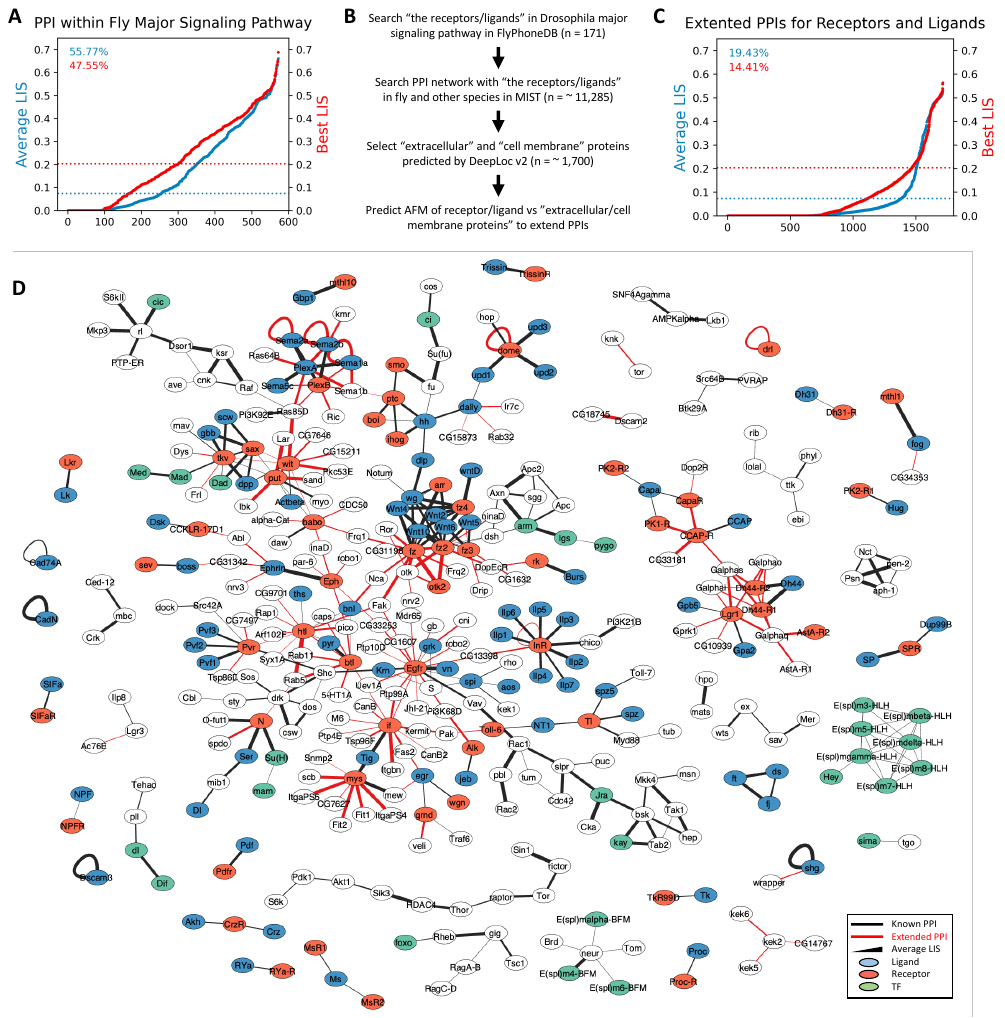

**Supplementary Figure 20. Expanding PPIs for Receptors and Ligands in *Drosophila* Major Signaling Pathways.**

A. Distribution of average and best LIS for components of the *Drosophila* Major Signaling Pathway (MSP) as annotated in FlyPhoneDB. The percentages above each graph represent the proportion of PPIs that exceeds the established LIS thresholds.
B. Strategy for predicting extended PPI sets involving MSP receptors and ligands. We compiled a list of 171 receptor and ligand proteins from these pathways as documented in FlyPhoneDB. Initial PPIs were extracted from the MIST database, which encompasses PPI data from *Drosophila* and various species, yielding 11,285 potential interactions. We narrowed our focus to approximately 1,700 PPIs involving predicted extracellular and membrane proteins based on DeepLoc v2 predictions. Further AFM analyses were then conducted to extend the PPI network, specifically seeking new interactions between MSP receptors/ligands and associated extracellular or membrane proteins.
C. Distribution of average and best LIS for newly predicted PPIs between MSP receptors/ligands and potential interactors, including extracellular and membrane proteins. The percentages at the top of each graph denote the fraction of PPIs that exceed each LIS thresholds.
D. A network representation of the extended *Drosophila* MSP, incorporating the additional PPIs. Existing PPIs from FlyPhoneDB are depicted with black edges, while newly predicted extensions for receptors and ligands are shown with red edges. The thickness of an edge correlates with the average LIS. The network nodes are color-coded: blue for ligands, orange for receptors, and green for transcription factors (TFs).

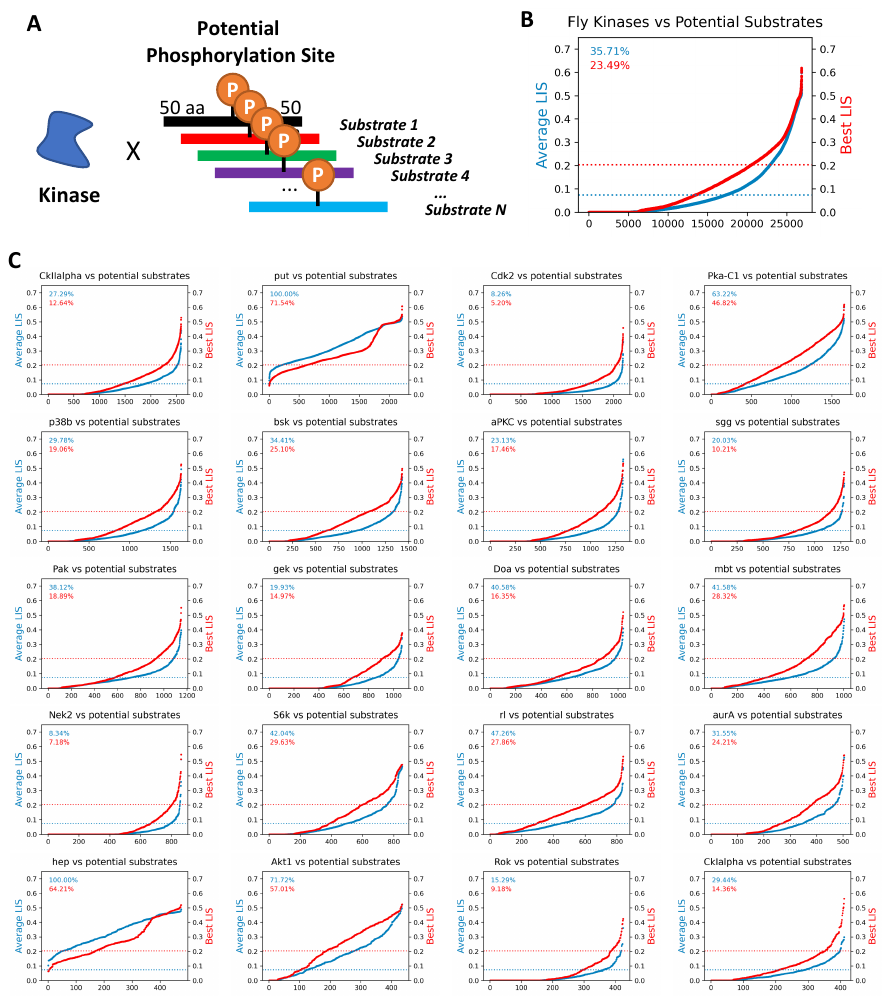

**Supplementary Figure 21. AFM screening for *Drosophila* kinases and their substrates.**

A. Schematic illustration of the AFM-based screening for potential kinase-substrate pairs. A total of 49 fly kinases were selected, with potential substrates defined by sequences extending 50 amino acids on either side of the phosphorylation sites.
B. Distribution of average and best LIS for 49 kinases and their potential substrates. The percentages above each graph represent the proportion of PPIs that exceed the established LIS thresholds.
C. Distribution of average and best LIS for top 20 kinases and their potential substrates. The percentages above each graph represent the proportion of PPIs that exceed the established LIS thresholds.

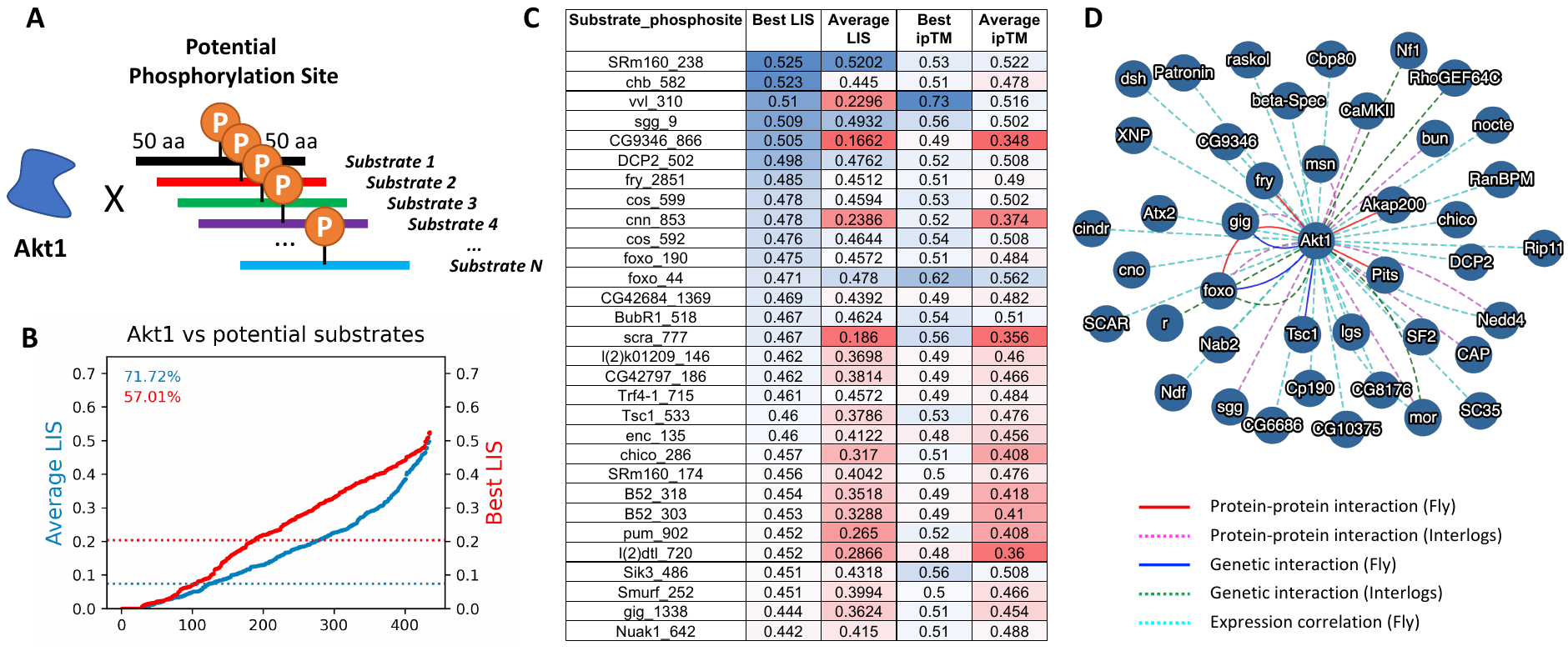

**Supplementary Figure 22. AFM screening for *Drosophila* Akt1 kinase and its substrates.**

A. Schematic illustration of the AFM-based screening method for Akt1 kinase and potential substrates, with each substrate represented by the amino acid sequence 50 residues upstream and downstream from the phosphorylation site.
B. Distribution of average and best LIS for Akt1 and its potential substrates. The percentages above each graph represent the proportion of PPIs that exceed the established LIS thresholds.
C. Table presenting the top 30 substrates ranked by the best LIS value, accompanied by their corresponding average LIS, best ipTM, and average ipTM scores.
D. Network visualization of Akt1-potential substrates with the interactions above the optimal best LIS threshold (0.201). The newtork was built in MIST. PPIs are shown by solid red lines for interactions within the fly species and magenta dotted lines for interactions inferred from interlogs. Genetic Interactions (GI) are illustrated with solid blue lines for interactions within the fly species and green dotted lines for interactions inferred from interlogs. Expression correlations between Akt1 and potential substrates, if present, are depicted with cyan dotted lines.
